# Temporal effects of a single oral dose of psilocybin on plasma circulating miRNAs in healthy young adults

**DOI:** 10.64898/2026.08.04.742716

**Authors:** Aedan O’Shea, Natasha Mason, Rudy Schreiber, Marcha C.T. Verheijen, Johannes G. Ramaekers, Jacco J. Briedé, Julian Krauskopf

**Affiliations:** Department of Translational Genomics, Faculty of Health, Medicine and Life Sciences, Maastricht University, Maastricht, The Netherlands; Department of Neuropsychology and Psychopharmacology, Faculty of Psychology and Neuroscience, Maastricht University, Maastricht, The Netherlands; Mental Health and Neuroscience Research Institute (MHeNs), Maastricht University, Maastricht, The Netherlands; Centre for integrative Neuroscience (CIN), Maastricht University, Maastricht, The Netherlands; GROW Institute of Oncology and Reproduction, Maastricht University, Maastricht, the Netherlands

## Abstract

**Background:** Psilocybin, a classic psychedelic, produces acute alterations in brain function, and has shown sustained therapeutic effects in psychiatric disorders. However, most studies focus on acute brain imaging readouts (e.g., fMRI) and rarely assess longer-term molecular changes. MicroRNAs (miRNAs) are small non-coding RNAs that regulate gene expression, are enriched in the brain, and can be released into blood, potentially indexing brain-relevant molecular processes. We hypothesised that a single psilocybin dose would show acute changes in miRNAs with predicted relevance to neuroplasticity related signalling and immune/inflammatory regulation in healthy adults.

**Methods:** In a randomised, double-blind, placebo-controlled study (N=62; 31 psilocybin, 31 placebo), volunteers received psilocybin (0.17 mg/kg) or placebo. Blood was collected at baseline, 360 minutes, and 7 days after dosing. Plasma miRNAs were quantified by small RNA sequencing. Elastic net regression was used for feature selection, followed by differential expression analysis and validation with linear mixed models. Pathway enrichment used Reactome and Gene Ontology.

**Results:** Two circulating miRNAs (let-7g-5p and miR-150-5p) met criteria for differentially expressed at 360 minutes following psilocybin administration, with no significant differences detected at 7 days or in the placebo condition under the statistical thresholds used. Over representation analysis suggested enrichment of molecular processes involved in neuroplasticity (e.g., TrkA, MAPK), inflammation (e.g., IL-6, TGF-β), and transcriptional regulation (e.g., RNA polymerase II, SMAD2/3/4).

**Conclusions:** A single oral dose of psilocybin was associated with transient alterations in circulating miRNA expression, consistent with an acute shift in circulating gene-regulatory miRNA signals, without sustained miRNA changes at 7 days. These findings provide initial evidence that circulating miRNA changes after psilocybin may reflect acute molecular process responses and support further investigation of circulating miRNAs as potential biomarkers of psychedelic-induced molecular responses.

## INTRODUCTION

Psilocybin and its pharmacologically active metabolite, psilocin (4-hydroxy-N,N-dimethyltryptamine), were identified as the compounds responsible for the psychedelic effects of psychedelic mushrooms in the 1950s ^1^. Both psilocybin and psilocin are derived from a tryptamine scaffold, similar to serotonin (5-hydroxytryptamine or 5-HT), making psilocybin a serotonergic classical psychedelic ^2,3^. Psilocin acts as a non-selective partial agonist of the serotonin-2A receptor (5-HT2AR), leading to profound subjective experiences, including changes in mood, cognition, and sensory perception ^2,4,5^.

A growing number of clinical trials suggest psilocybin may provide therapeutic relief for a range of psychiatric disorders, including anxiety, Substance-Use Disorder (SUD), and Major Depressive Disorder (MDD), particularly Treatment-Resistant Depression (TRD) ^6–9^. In some studies, therapeutic effects persist long after acute drug exposure, and it has been hypothesized that these sustained effects might involve increased neuroplasticity and/or decreased inflammation.

Research on the biological effects of psilocybin in humans has largely focused on acute brain activity and functional connectivity changes measured using fMRI. Notably, longitudinal precision functional mapping suggests that psilocybin exposure is associated with persistent functional connectivity alterations lasting for weeks ^10^. Mechanistically, psilocin’s actions at the 5-HT2AR, among other targets, are thought to be key mediators of psychedelic effects and are proposed to initiate plasticity-relevant cascades involving enhanced glutamatergic signalling and synaptic remodelling ^11–13^. These cascades may engage canonical intracellular signalling, including calcium/calmodulin-dependent protein kinase II (CaMKII) and mitogen-activated protein kinase/extracellular signal-regulated kinase (MAPK/ERK) pathways, which regulate activity-dependent gene expression relevant to synaptic plasticity ^5^. Activation of downstream targets, including brain-derived neurotrophic factor (BDNF) signalling via tropomyosin receptor kinase B (TrkB), mammalian target of rapamycin (mTOR) signalling, and AMPA-type glutamate receptor engagement, may also be necessary to produce sustained neuroplastic effects (Jaster et al., 2022; Olson, 2022). Despite this mechanistic rationale, peripheral BDNF measures yield inconsistent findings in humans following psychedelic administration, likely attributable to measurement-related variability, limiting their utility as a standalone biomarker of plasticity-related signalling ^14–16^. Consequently, there is an unmet need for minimally invasive biomarkers that can reflect downstream molecular processes relevant to neuroplasticity in humans.

Beyond neuroplasticity, psilocybin has also been reported to exhibit anti-inflammatory properties. Patients with stress-related psychiatric disorders such as MDD often have elevated levels of inflammatory markers, including Tumor Necrosis Factor-alpha (TNF-α), Interleukin-6 (IL-6), and C-reactive protein (CRP) ^17,18^. In a previous clinical study, psilocybin showed acute and persisting effects on circulating inflammatory markers. A single psilocybin dose acutely reduced TNF-α levels, which returned to baseline within seven days, while IL-6 and CRP remained unchanged acutely, but were reduced at seven days, indicating a sustained anti-inflammatory effect ^19^. To date, the molecular mechanisms underlying psilocybin’s sustained neuroplastic and anti-inflammatory effects remain poorly understood.

To better characterise the temporal molecular effects of psilocybin in humans, longitudinal profiling of blood-based gene-regulatory signals may provide insight into downstream molecular processes relevant to neuroplasticity and inflammation. MicroRNAs (miRNAs) are single-stranded, non-coding RNAs typically 18-25 nucleotides in length that act as significant regulators of gene expression at the post-transcriptional level. They act by binding to complementary sequences in the 3’-untranslated region of target messenger-RNAs (mRNAs), primarily through interactions with the miRNA seed sequence ^20^. This binding typically causes mRNA destabilization or translational repression, leading to gene silencing ^21^. Each miRNA can regulate multiple mRNA targets, influencing a wide range of cellular processes. Importantly, miRNAs play critical regulatory roles in the brain and can be secreted or released into the bloodstream, often via extracellular vesicles (EVs) such as exosomes, which are capable of crossing the blood–brain barrier (BBB) and thereby reflecting molecular changes in the nervous system ^22–24^. When miRNAs are present in bodily fluids including blood plasma, they are called circulating miRNAs (cimiRNAs) and can be obtained through minimally invasive methods. Therefore, analysing cimiRNA expression profiles may offer a window into the molecular processes underlying psilocybin’s effects on the nervous system ^20,21^.

The present study employs an exploratory approach to assess how a single low-to-moderate oral dose of psilocybin in healthy human volunteers may influence neuroplasticity and inflammation-related biological processes, as measured by changes in cimiRNA expression profiles. The cimiRNAs identified to be Differentially Expressed (DE) in this study may provide preliminary insight into the molecular pharmacodynamics of psilocybin. This analysis may inform future research exploring Differentially Expressed miRNAs (DEmiRs) as potential biomarkers to monitor treatment response and guide targeted interventions in psychiatry, due to the elusive mechanism of action underpinning psilocybin’s potential therapeutic benefits.

## MATERIALS & METHODS

### Participants and study design

Plasma circulating miRNA expression profiles were assessed in the blood plasma of 62 healthy adult participants as part of a randomised, placebo-controlled, double-blind, parallel-group design ^4^. Participants were allocated to treatment groups and received either a single low-to-moderate oral dose of psilocybin (0.17 mg/kg; n=31) or placebo (n=31). All participants had previous experience with psychedelic drug usage. Exclusion criteria included: psychedelic drug use <3 months; history of drug abuse or addiction; pregnancy or lactation; health issues including hypertension (diastolic >90 and systolic >140); cardiac dysfunction; liver dysfunction; current or history of psychiatric disorders and previous experience of serious side effects from cannabis. A study design scheme depicting the parallel group design is displayed in *Figure 1.* The treatment was administered orally, in a closed cup containing bitter lemon (placebo) or bitter lemon with psilocybin (powder). Treatment groups were matched for age, sex, and education level (*Table 1*). This study was conducted according to the code of ethics on human experimentation established by the declaration of Helsinki (1964) and amended in Fortaleza (Brazil, October 2013) and in accordance with the Medical Research Involving Human Subjects Act (WMO) and was approved by the Academic Hospital and University’s Medical Ethics committee (PMID: 32446245). All participants were fully informed of all procedures, possible adverse reactions, legal rights and responsibilities, expected benefits, and their right for voluntary termination without consequences. Written informed consent was obtained from all participants.

**Figure 1.**
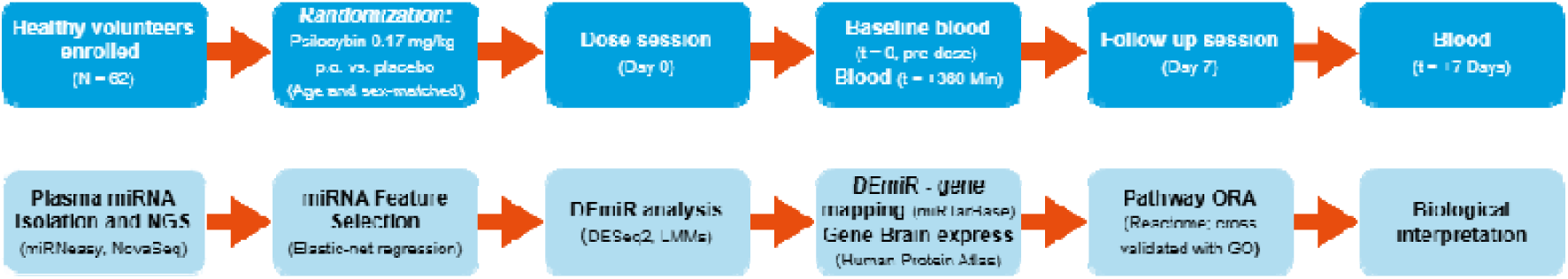
Study design. Sixty-two healthy volunteers were randomized to receive a single p.o. dose of psilocybin (0.17 mg/kg) or placebo. Blood plasma samples were collected at baseline (pre-dose), 360 minutes, and 7 days post-dose for circulating miRNA profiling. MiRNAs were isolated and sequenced by Next-Generation Sequencing (NGS) and differentially expressed miRNAs (DEmiRs) were identified using DESeq2 and linear mixed models following elastic net based feature selection. DEmiRs were mapped to their target genes (miRTarBase) and restricted to brain-expressed genes (Human Protein Atlas). Gene-set over-representation analysis was then performed using Reactome, with pathways cross validated using Gene Ontology to highlight biological processes enriched for DEmiR target genes.

**Table 1:**
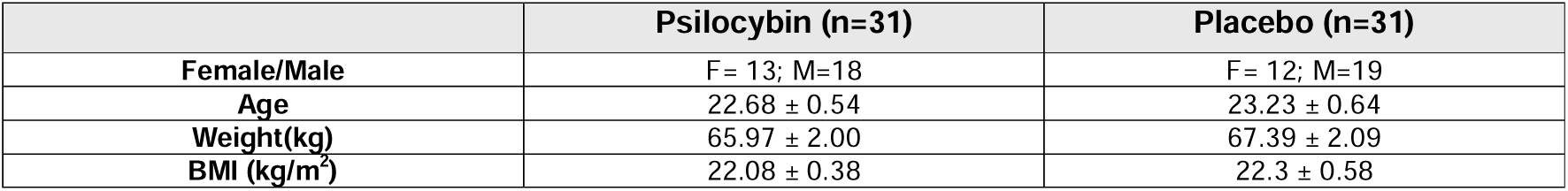
demographic characteristics of the participants; Females/Males per group, age (mean ± SEM), weight in kilograms (mean ± SEM), and body mass index (BMI) (mean ± SEM).

### Demographics

A total of 62 adult participants (37 male, 25 female), age 18-34 years, participated in the current study. A net total of 170 plasma samples were obtained. Table 1 presents the demographic characteristics of the participants, including mean weight, and body mass index (BMI).

### Blood sample processing, miRNA isolation and Next generation sequencing (NGS)

Blood samples were collected in K3-EDTA tubes (Vacutest Kima, 13010) at baseline, 360 minutes, and 7 days post-administration. Blood plasma was obtained following immediate centrifugation at 3500 x g in 4°C for 10 minutes, before being stored at −80 °C prior to analysis. Haemolysis was measured in each sample using a Nanodrop 1000 (Life Technologies) at a wavelength of 414nm ^25^. Haemolytic samples (N= 10) were excluded from the study. Total Small-RNAs, including miRNAs, were extracted from 200μl of plasma using the miRNeasy Serum/Plasma Kit (Qiagen) following the manufacturer’s protocol. The isolated RNA was eluted in 14μl of RNase-free water. Small RNA libraries were generated using the NEXTFLEX Small RNA-seq Kit v3 (PerkinElmer, catalog #NOVA-5132) from 4.75 μl of RNA as input with the optional tRNA/yRNA blocker and unique dual indexes (UDIs). A no-size-selection approach was applied, as recommended for cell-free RNA. The protocol included 3’ and 5’ 4N adenylated adapter ligation, followed by excess adapter removal with NEXTFLEX clean-up beads. UDI barcodes were incorporated during ligation, and complete libraries were amplified using 20 PCR cycles. Library quantification was performed using the Qubit HS dsDNA kit on a Qubit fluorometer 2.0 (Life Technologies), and fragment size distribution was assessed using High Sensitivity D1000 tapes on an Agilent TapeStation 2200. Final libraries were normalized to a concentration of 1.6 nM, pooled at 2 μl per sample, and sequenced using a 100-cycle SP flowcell on the Illumina NovaSeq 6000 with a 1×100 bp read length. A total of 971.3 million sequencing reads were obtained, with an average of 5.49 million reads per sample (median: 5.4 million).

### Data processing and Differential Expression analysis

Plasma small RNAs were profiled by Next-Generation Sequencing (NGS) without target preselection, and raw sequencing data were de-multiplexed, and FASTQ files were generated using bclfastq (version 2.20.0) (Illumina). Adapter and 4N sequence trimming were performed using Cutadapt (version 3.1), and miRNA quantification was carried out with miRge3.0 ^26^. Reads were mapped to the human miRge3 library using Bowtie (version 1.3.1) with default parameters. Small RNA-sequencing data have been deposited in NCBI GEO under accession GSE319559.

DE analysis was performed using the data filtering recommendations of the R-ODAF pipeline ^27^. First, 10 haemolytic samples were excluded. Also, a coverage threshold of 1,000,000 total mapped reads per sample was applied, excluding 7 samples from the analysis. Principal component analysis (PCA) was used to identify outlier samples, with >20% variance between samples used as a threshold for exclusion, resulting in the exclusion of 3 samples. In total, 150 samples passed the filtering criteria. A full overview of the sample exclusions can be viewed in *Table 2*.

**Table 2:** CONSORT style sample flow showing collected samples, exclusions due to haemolysis, low sequencing depth, and PCA outliers, stratified by treatment group and timepoint.

|  | Total | Psilocybin | Placebo | Baseline | +360 Min | +7 Days | Reason |
| --- | --- | --- | --- | --- | --- | --- | --- |
| Possible samples | 186 | 93 | 93 | 62 | 62 | 62 | Design maximum |
| Collected samples | 170 | 86 | 84 | 59 | 57 | 54 | Missing blood samples |
| Passed QC | 150 | 75 | 75 | 54 | 49 | 47 | Final analysis set |
| Excluded: haemolysis | 10 | 5 | 5 | 2 | 5 | 3 | Haemolysis |
| Excluded: sequencing depth | 7 | 4 | 3 | 2 | 3 | 2 | Low reads |
| Excluded: PCA outlier | 3 | 2 | 1 | 1 | 0 | 2 | Outlier |

From the total of 2,731 miRNAs in the refence genome, 500 miRNAs were consistently expressed (>80% of samples in one of the treatment groups). Thereafter, a two-pass analysis approach was employed to identify psilocybin exposure-relevant cimiRNAs. To identify miRNAs associated with psilocybin exposure, elastic net regression (α = 0.05) with a binomial logistic approach was applied to perform miRNA ‘feature selection’, using the glmnet package (version 4.1.8) in R. MiRNA expression data were used as predictors, and psilocybin exposure status (treatment vs. placebo) was the outcome. The optimal penalty parameter (λ) was selected through cross-validation, and miRNAs with non-zero regression coefficients were retained. The mixing parameter (α = 0.05) emphasized the ridge component to account for correlations amongst miRNAs, while incorporating limiter sparsity from the elastic net penalty. This reduced overfitting and stabilized coefficient estimates ^28^. Following the filtering steps described, 56 miRNAs were used for further analysis. These same 56 psilocybin-associated miRNAs were then used to test DE under the placebo condition.

Finally, miRNAs were assessed through DE analysis, using the DESeq2 (version 1.44.0) in R (version 4.4.1). Analyses were performed separately within psilocybin and placebo groups. Comparisons made were baseline vs. +360 minutes post psilocybin exposure, and baseline vs. +7 Day follow up. Batch effect, Age, Sex and BMI were each found to be significant confounding variables and therefore included in the design of DEseq2 to minimize confounding (Supplementary 9). DEmiRs identified using DESeq2 with a False Discovery Rate (FDR) <0.05 were deemed to be associated with psilocybin exposure.

To ensure statistical robustness, selected psilocybin-associated miRNAs were re-evaluated using linear mixed-effects models (LMMs) implemented in lmerSeq (version 0.1.7) ^29^. For each miRNA, variance-stabilized expression, which provides robust handling of correlated, longitudinal RNA-sequencing data, was modelled as the outcome and timepoint (baseline, 360 min, or 7 days) as the primary exposure, adjusting for sex and haemolysis, with random intercepts for batch and subject ID. This framework accounted for technical variation and parallel-group correlation.

Hypothesis testing focused on changes at +360 min vs. baseline and +7 days vs. baseline. P-values were adjusted using the Benjamini–Hochberg procedure, and associations with FDR < 0.10 were considered significant. Models were fitted separately for psilocybin and placebo groups. An FDR threshold of 0.10 was used for the LMMs sensitivity analyses because this study was exploratory, modestly powered, and designed to identify candidate circulating miRNA signals for future validation rather than to establish definitive biomarkers. This threshold was selected to balance false-discovery control with sensitivity in the context of a first-in-human peripheral miRNA profiling study of psilocybin.

### Pathway analysis and biological interpretation

To establish how DE miRNAs may elicit an effect on their target genes, experimentally validated target genes of DE miRNAs were obtained from miRTarBase (version 9.0). Functional miRNA-Target Interactions (MTI) with strong evidence (e.g., Luciferase reporter assays, Western blot, RT-PCR, or CLIP-based methods) were used. Tissue specificity of the DE miRNAs and their target genes was evaluated using miRNATissueAtlas2 and The Human Protein Atlas (version 24.0) respectively ^30^. Genes which are known to be expressed in the brain were considered, specifically in the following regions: amygdala, basal ganglia, cerebellum, cerebral cortex, choroid plexus, hippocampal formation, hypothalamus and midbrain. Over-representation analysis (ORA) was performed using the Reactome database (https://reactome.org, accessed 08/04/2025). For each DEmiR, ORA was carried out using its experimentally validated target genes, filtered for brain expression, as input to test whether Reactome pathways contained more of these genes than would be expected by chance. Pathways with ≥2 mapped genes and an FDR < 0.05 were deemed to be significantly overrepresented. Alongside the full Reactome output, a summarized version of the results was used. Summarization was based on the hierarchical structure of the database. For each overrepresented pathway, the main biological process and the first sub-pathway in the hierarchy were manually identified. A list of the Reactome sub-pathways and their prevalence is displayed in *Supplementary Tables 1 and 5*.

Additionally, Gene Ontology (GO) ORA was utilized to support the results achieved from Reactome ORA. We cross-validated Reactome pathway results with GO term enrichment to assess consistency. Both Reactome and GO outputs were filtered for significance (FDR ≤ 0.05) and converted into gene to term lists. Terms from Reactome and GO were then matched when they shared ≥2 genes.

This integrated approach allowed for prioritization of biologically meaningful processes potentially regulated by each DEmiR in a brain directed context. Once biologically relevant pathways were identified, their influence on processes related to neuroplasticity and inflammation were investigated through a manual literature review. This was performed using the keywords “neuroplasticity”, “neurogenesis”, “synaptogenesis”, “spinogenesis”, “synaptic remodelling”, “synapse,” “axon,” “neuronal,” “memory,” “learning”, “inflammation”, “immune,” “immune signalling “, “cytokine,” “tumor necrosis,” “interleukin,” and “chemokine”. This step was used to contextualize the functional significance of the enriched pathways within brain-related processes.

### EXOmotif

To investigate sequence features that may influence the extracellular release of miRNAs, we performed a motif-based analysis focused on exosomal motifs (EXOmotifs). The EXOmotif sequences were obtained from Garcia-Martin et al. who reported short nucleotide motifs that mediate selective miRNA loading into exosomes and small-EVs ^31^. Identified EXOmotifs included, among others, the CAUG and GAGG elements. All candidate miRNA sequences in our dataset were scanned for the presence of these motifs in both the 5′ to 3′ and reverse complement orientations to ensure comprehensive detection. For each miRNA, the number and type of EXOmotifs were recorded, allowing classification of miRNAs according to their predicted export into exosomes.

## RESULTS

### Differential Expression Analysis of Circulating-miRNAs

To assess psilocybin-associated changes in circulating miRNA expression, we first performed DE analysis using DESeq2. The comparisons made by our analysis were +360 minutes post psilocybin exposure vs. Baseline, and +7 Day follow up vs. Baseline. Batch, age, sex, and BMI were included as covariates. In the +360 minutes post psilocybin exposure vs. Baseline comparison by our DESeq2 analysis, five cimiRNAs were found to be DE (FDR<0.05). While no DEmiRs were found at the +7 Day post-psilocybin follow up, nor in either of the placebo conditions. Of the five DEmiRs identified by DESeq2 in the +360 minutes vs. Baseline comparison, three were upregulated while two were downregulated +360 minutes post psilocybin exposure, as can be seen by Log2FoldChange (L2FC) shown in *Table 3*.

**Table 3:**
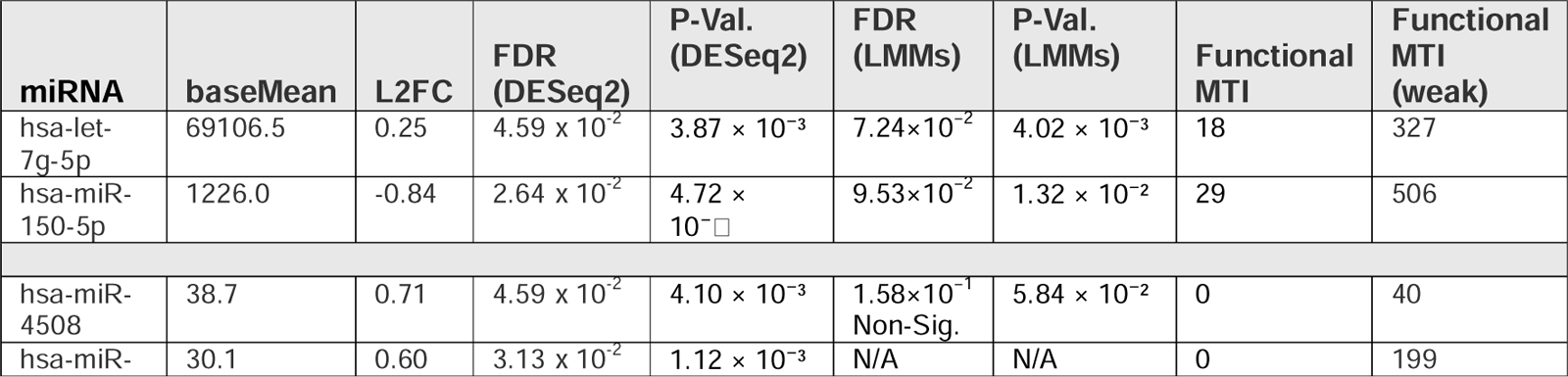
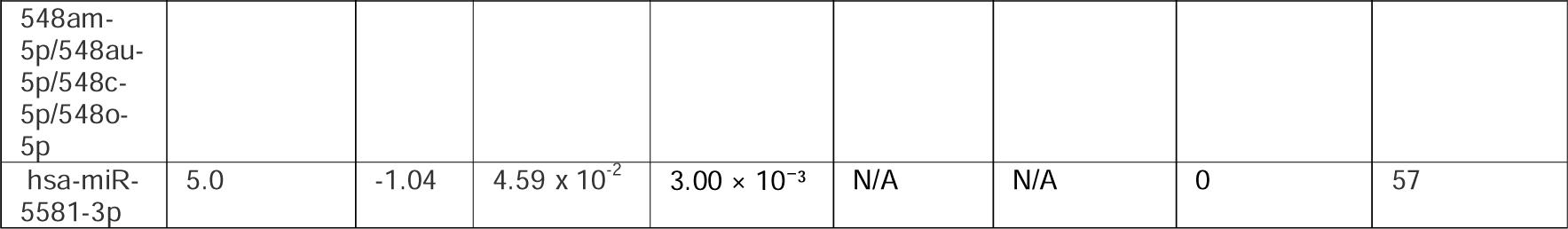
Table displaying DEs miRNAs associated with psilocybin exposure identified by +360 minutes post psilocybin vs. Baseline comparison. DEmiRs listed with their Mean read count (baseMean), Log2FoldChange (L2FC), and nominal P-values alongside P-adjusted value by Benjamini-Hochberg procedure (FDR) achieved from both DESeq2 and LMMs analysis. Positive L2FC values indicate higher expression relative to baseline while negative L2FC values indicate lower expression relative to baseline. Number of target genes using validated interactions from miRTarBase at different validations levels {Functional MTI / Functional MTI (weak)}.

To ensure statistical robustness of the five DEmiRs identified by DE analysis, the initial feature selected miRNAs were re-evaluated using LMMs. Variance-stabilized expression was modelled as the outcome, with timepoint as the primary exposure, adjusting for sex and haemolysis, and random intercepts for batch and subject ID. P-values were adjusted using the Benjamini–Hochberg procedure (FDR <0.10). Following LMMs validation, ‘hsa-let-7g-5p’ and ‘hsa-miR-150-5p’ remained significantly associated with psilocybin exposure at 360 minutes (α=0.10). Each of these DEmiRs also had stronger experimental evidence of miR targets, labelled as Functional MTI, reported in miRTarBase (version 9.0) shown in *Table 3.* Both DEmiRs later returned to baseline and thus no differences were observed at the 7-day follow-up. These patterns are illustrated in *Figure 2*.

**Figure 2:**
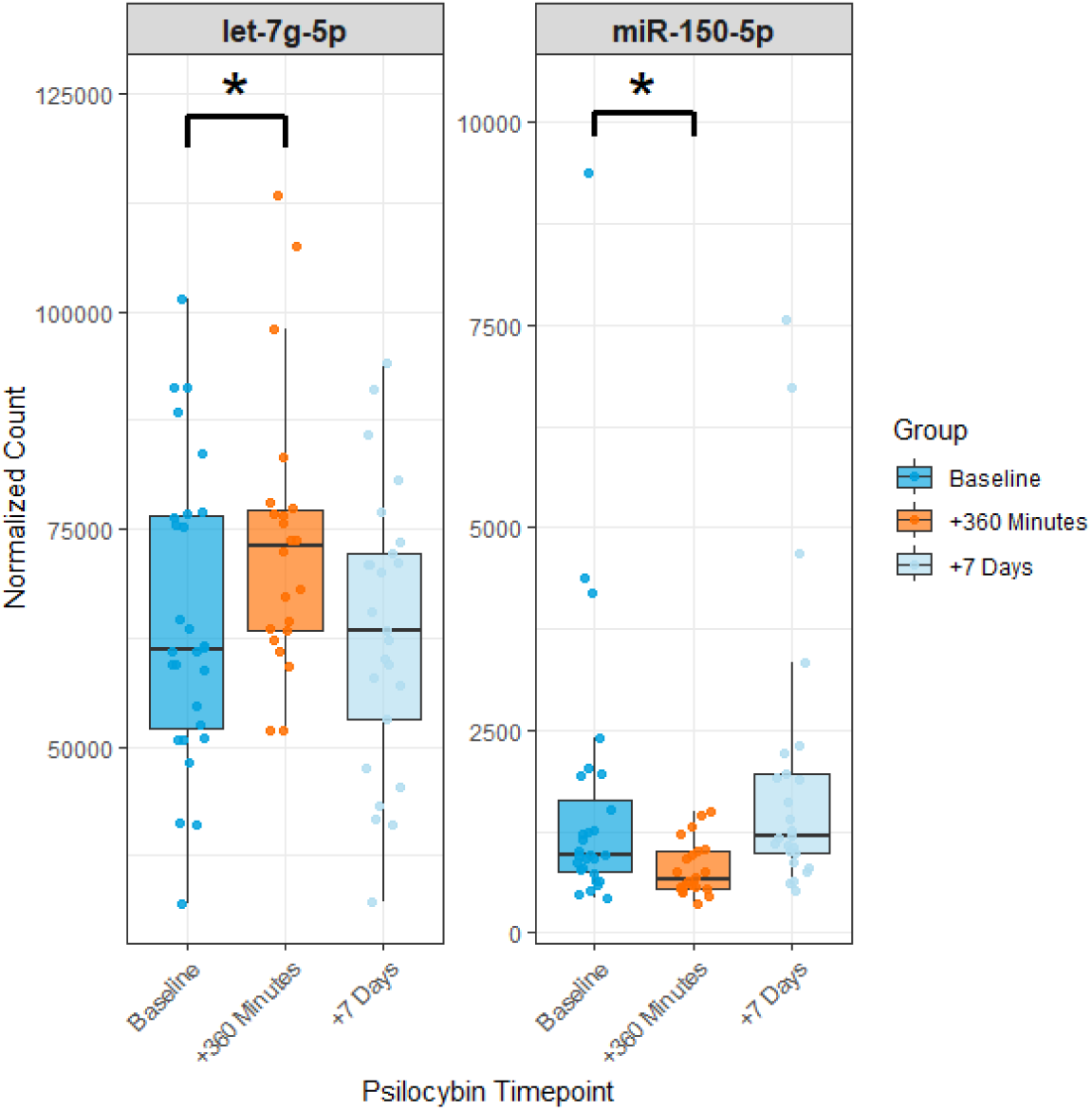
Boxplots of normalized read counts for differentially expressed miRNAs identified by both DESeq2 and LMMs, +360 minutes post psilocybin exposure. Expression levels are shown at baseline, 360 minutes, and 7 days post-exposure. Dark blue boxes indicate baseline, orange boxes indicate 360 minutes, and light blue boxes indicate 7 days. Individual points represent sample-level normalized counts. Brackets with asterisks denote statistically significant differences between baseline and 360 minutes (DESeq2 FDR-adjusted p < 0.05, LMMs FDR-adjusted p < 0.10).

### Pathway analysis and biological interpretation

We used miRTarBase to identify miRNA-Target Interactions (MTIs) for the two DEmiRs identified. Both hsa-let-7g-5p and hsa-miR-150-5p had strong experimental support based on validation by functional MTIs (e.g., luciferase reporter assays, Western blot, RT-PCR, or CLIP-based methods), with evidence indicating their regulation of 18 and 29 target genes respectively.

After MTIs were identified, target genes were filtered to include only those which are expressed in the brain, via The Human Protein Atlas (version 24.0). The brain regions used in this filtering step were: “amygdala”, “basal ganglia”, “cerebellum”, “cerebral cortex”, “choroid plexus”, “hippocampal formation”, “hypothalamus” and “midbrain”. A total of 3 genes modulated by our chosen DEmiRs were not expressed in the brain and subsequently removed. The genes removed were MUC4 and NANOG, which are regulated by hsa-miR-150-5p, and IGF2BP1, which is target of hsa-let-7g-5p.

### Over Representation Analysis of DEmiRs: FDR >0.05 and +/- 2 genes

To gain insight to the cellular processes effected by psilocybin exposure, experimentally validated gene targets of the identified DEmiRs were used to perform pathway enrichment analysis via ORA using the Reactome database, which was then supported by ORA using GO. A list of included genes can be found in Supplementary 10.

### Let-7g-5p

The upregulated DEmiR ‘let-7g-5p’ was found to regulate 17 target genes expressed in the brain. Reactome ORA using this set of genes yielded a total of 99 significant pathways (FDR <0.05, ≥2 genes), while GO ORA yielded a total of 244 significant terms (FDR <0.05, ≥2 genes). For example, target genes of let-7g-5p are implicated in ‘Generic Transcription Pathway’ in Reactome, and ‘Cellular response to abiotic stimulus’ in GO. The 10 most significantly over-represented pathways derived from Reactome ORA and GO ORA are displayed below in *Figure 3*.

**Figure 3:**
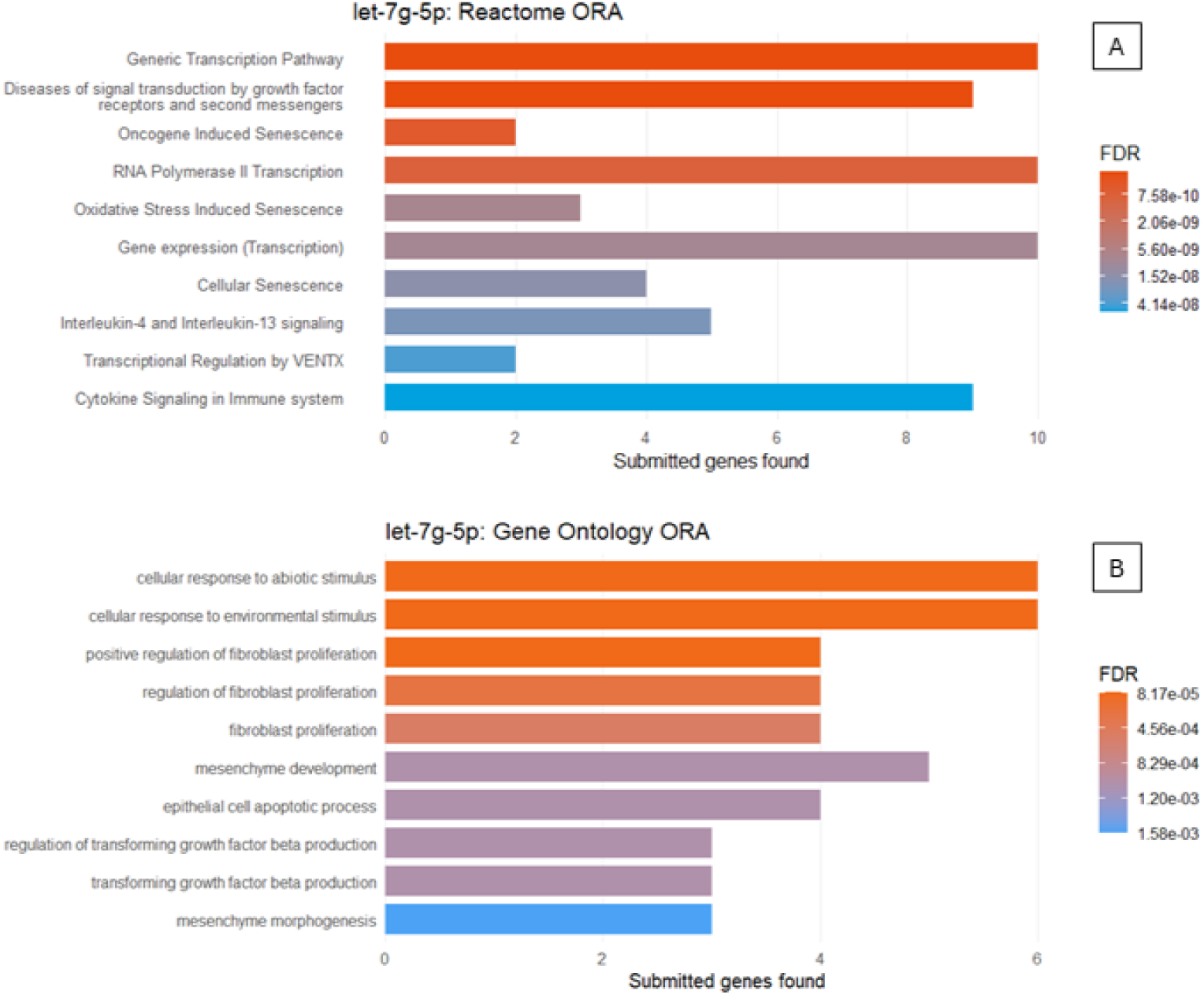
10 most significantly Over-represented pathways identified by Reactome ORA (A) and GO ORA (B) for gene-sets of brain-expressed genes experimentally validated to be targets of the DEmiR let-7g-5p. Horizontal bars indicate the number of submitted gene targets per pathway (X-axis), colour represents FDR-adjusted significance.

Comparison of Reactome and GO ORA for let-7g-5p by gene overlap yielded 16 unique Reactome pathways, and 192 unique GO terms with ≥2 genes overlapping respectively. A list of the Reactome pathways, including information regarding the number of cross-validated GO terms can be viewed in *Supplementary Tables 2-4*.

### miR-150-5p

The downregulated DEmiR ‘miR-150-5p’ was found to regulate 27 target genes expressed in the brain. Reactome ORA yielded a total of 56 significant pathways (FDR <0.05, ≥2 genes). For example, target genes of miR-150-5p are implicated in ‘Signaling by Interleukins’ in Reactome, and ‘Response to hypoxia’ in GO.

The 10 most significantly over-represented pathways derived from Reactome ORA and GO ORA are displayed below in *Figure 4*.

**Figure 4:**
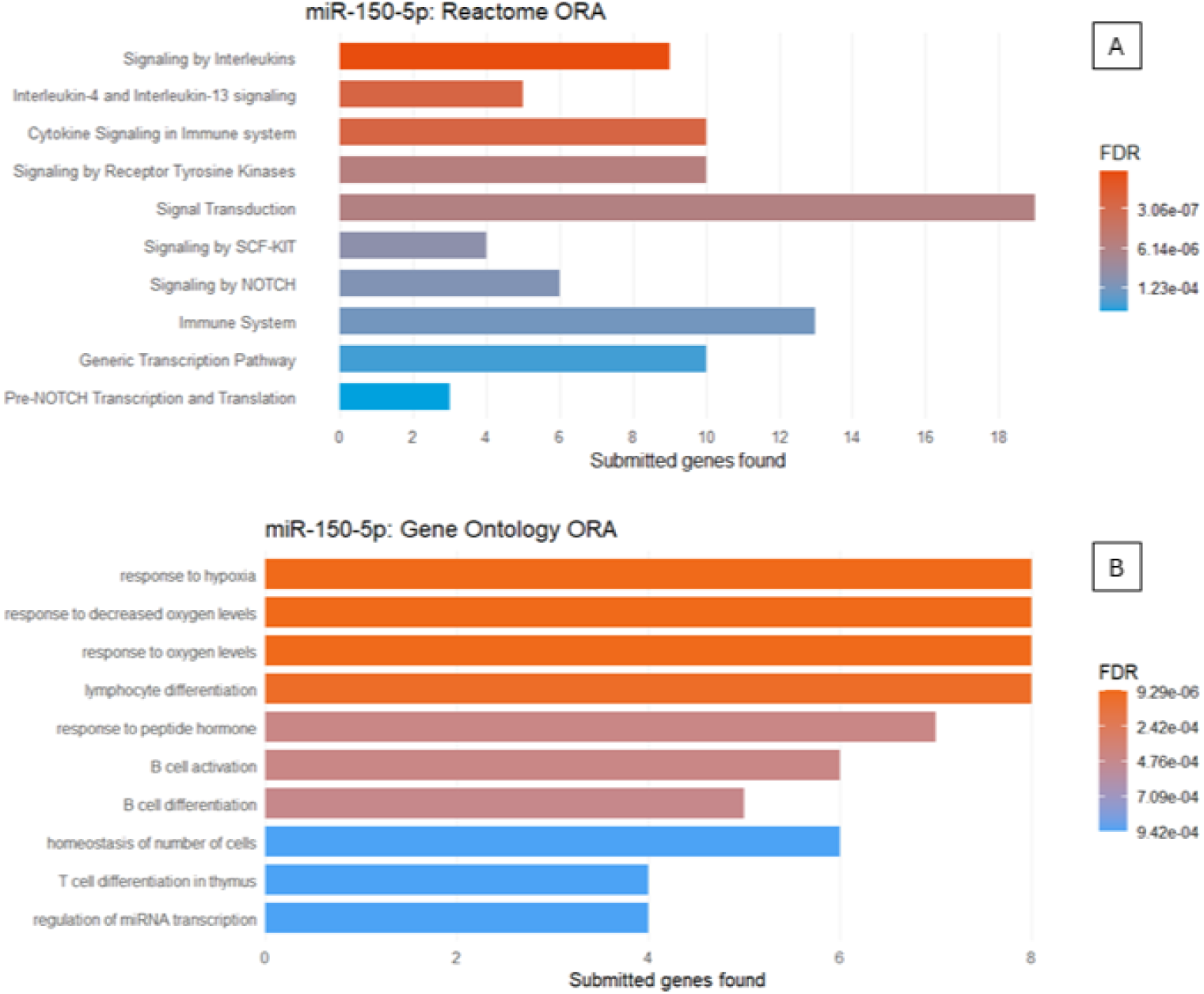
10 most significantly Over-represented pathways identified by Reactome ORA (A) and GO ORA (B) for gene-sets of brain-expressed genes experimentally validated to be targets of the DEmiR miR-150-5p. Horizontal bars indicate the number of gene targets per pathway (X-axis), colour represents FDR-adjusted significance.

Comparison of Reactome and GO ORA for miR-150-5p by gene overlap yielded 17 unique Reactome pathways, and 246 unique GO terms with ≥2 genes overlapping respectively. A list of the Reactome pathways, including information regarding the number of cross-validated GO terms can be viewed in *Supplementary Tables 6-8*.

Overall, a total of 24 unique Reactome pathways were over-represented and verified by supporting GO terms with ≥2 common genes. Of which, 9 were shared between both DEmiRs (Supplementary 8). The DEmiRs identified at the +360minutes time point in our analysis appear to modulate varying biological processes, reflected by overrepresented Reactome pathways associated with inflammation and neuroplasticity. Furthermore, over-represented transcription pathways may facilitate a sustained effect onset by psilocybin exposure.

### EXOmotif Identification

In our investigation of sequence features that may influence the extracellular release of miRNAs, Motif analysis of the selected miRNAs revealed that hsa-miR-150-5p contained a single exosomal motif (CAUG), whereas hsa-let-7g-5p contained two distinct exosomal motifs (CAUG and GAGG). No cell retention motifs were identified in either of the identified DEmiRs. These short sequence elements, previously described by Garcia-Martin and colleagues as EXOmotifs that guide miRNA sorting into exosomes, were clearly enriched in the analysed sequences ^31^.

## DISCUSSION

This study shows that a single low-to-moderate oral dose of psilocybin (0.17 mg/kg) is associated with a transient change in circulating miRNA expression in healthy adults. Two cimiRNAs were differentially expressed at +360 minutes, which then returned to baseline levels at the +7 day follow up. This suggests an acute but time-limited influence of psilocybin exposure on miRNA expression. The DEmiRs, let-7g-5p and miR-150-5p showed differential expression in opposing directions, as ‘let-7g-5p’ was found to be upregulated (L2FC; 0.25) while ‘mir-150-5p’ was downregulated (L2FC; −0.84). To our knowledge, this is the first randomized, placebo-controlled study reporting changes in the plasma circulating let-7g-5p and mir-150-5p miRNAs following psilocybin exposure in humans. The goal of this exploratory analysis was to map DEmiRs to candidate pathways for hypothesis generation, not to claim causality or therapeutic mechanism. More broadly, these findings support the use of circulating miRNAs as peripheral molecular readouts within emerging biomarker frameworks for rapid acting antidepressant and psychedelic interventions.

The two psilocybin-associated DEmiRs, let-7g-5p and miR-150-5p, identified by our analysis are predicted to modulate biological processes involved in neuroplasticity and growth factor signalling. The therapeutic effects of psychedelics like psilocybin, characterised by rapid onset and sustained duration, are hypothesised to result from increased neuroplasticity in brain regions such as the prefrontal cortex (PFC) and hippocampus ^5,32^. This interpretation is consistent with recent antidepressant literature, which proposes that psychedelics, ketamine and esketamine may act through enhanced brain plasticity despite distinct primary pharmacological targets ^33^. Prior research in rodents shows that psilocybin or psilocin can rapidly increase neuritogenesis, spinogenesis, and synaptogenesis in cortical neurons, and can induce region specific modulation of neuroplasticity associated genes in the prefrontal cortex and hippocampus ^34–36^. In the present dataset, target genes of these DEmiRs with experimentally validated miRNA–target evidence map to signalling cascades downstream of neurotrophin receptors, growth factor receptors, and MAPK pathways ^37^. The aforementioned processes align with preclinical evidence of psychedelic-induced plasticity, including TrkB- and BDNF-dependent signalling that have been implicated in structural remodelling and activity-dependent synaptic strengthening ^35,36,38,39^. We did not assess synaptic plasticity or brain tissue directly, so these pathway-level associations should not be interpreted as evidence that psilocybin induces neuroplasticity in humans. Instead, they suggest that circulating miRNAs may provide accessible peripheral readouts of acute signalling cascades associated with psilocybin and represent candidate blood-based biomarkers of acute molecular response.

The two DEmiRs identified were also linked to inflammatory and growth factor signalling. Reactome ORA was consistent with enrichment of cytokine pathways involving Interleukin-4, Interleukin-13, IL-6 and Transforming Growth Factor Beta (TGF-β), which regulate immune responses and influence neuronal and glial survival and tissue remodelling ^40,41^. This inflammatory component is relevant to the concept of immuno-metabolic depression (IMD), a proposed depression subtype characterized by low-grade inflammation, metabolic dysregulation and atypical energy-related symptoms, estimated to affect approximately 20–30% of individuals with depression ^42^. In a clinical study from which our samples were obtained, circulating IL-6 protein levels were significantly reduced 7 days after psilocybin but not after 360 minutes ^19^, while ORA in the present study showed enrichment of IL-6 signalling at 360 minutes. However, the IL-6 Reactome pathway was not supported by GO due to limited gene target overlap. This pattern is consistent with miRNA-level regulatory changes occurring at 360 minutes that may precede the delayed reduction in plasma IL-6 protein concentration observed at the later 7-day time point; however, this temporal relationship does not establish causality. Gene-sets of our DEmiRs also mapped to Vascular Endothelial Growth Factor (VEGF) and Platelet-Derived Growth Factor (PDGF) related pathways, which coordinate angiogenesis, cell survival, and synaptic and dendritic spine remodelling, and converge on MAPK signalling implicated in plasticity-related processes ^43–46^. In an interplay with BDNF, Nerve Growth Factor (NGF) has significant influence in neuroplasticity related processes including synaptic plasticity and dendrite development ^39,47^. The NGF-stimulated transcription pathway was also over-represented, which is known to cause cell differentiation and neurite outgrowth ^48^. Together, these associations suggest that psilocybin transiently engages coordinated immune and growth factor signalling at the pathway level. This does not imply that psilocybin directly treats IMD, because the present cohort consisted of healthy adults, this overlap should be interpreted as hypothesis generation rather than as evidence for relevance to IMD specifically.

Transcriptional regulation pathways were also over-represented. RNA polymerase II transcription was found to be the most significant transcription pathway by Reactome ORA for both DEmiRs, suggesting broad engagement of gene expression control. Enriched networks involving TP53, RUNX family transcription factors and SMAD-mediated signalling link growth factor and cytokine inputs, including TGF-β, to stress-responsive and plasticity-related transcriptional programmes, indicating intracellular immune-mediated crosstalk ^48–50^. This profile is consistent with a brief, coordinated adjustment of transcriptional control downstream of the observed miRNA shifts. Neurotrophic signalling, inflammatory signalling, and transcriptional reprogramming form a tightly connected regulatory loop that can shape neuronal structure, glial state, and cellular stress responses ^51^. Such multi-domain signalling is compatible with recent mechanism-driven models of TRD, which emphasize that rapid-acting interventions may act beyond monoaminergic neurotransmission by engaging glutamatergic, neuroplastic, inflammatory and neuroendocrine systems simultaneously ^33,52^.

We also explored whether the two DEmiRs carry sequence features associated with EV export. Motif analysis of the selected miRNAs revealed that hsa-miR-150-5p contained a single EXOmotif (CAUG), whereas hsa-let-7g-5p contained two distinct EXOmotifs (CAUG and GAGG). No cell retention motifs were identified in either of the identified DEmiRs. These short sequence elements, previously described by Garcia-Martin and colleagues were clearly enriched in the analysed sequences and correlated with miRNA sorting into exosomes ^31^. This pattern is compatible with selective packaging for intercellular transfer and suggests that psilocybin briefly alters the regulatory miRNA cargo available for systemic signalling, including towards the brain as vesicle mediated transport across BBB has been documented ^21–24^.

Both psilocybin associated DEmiRs identified in the present study have been implicated in neurodegenerative and affective disorder biology. Circulating let-7g-5p is reported as reduced and miR-150-5p as increased in several Alzheimer’s disease (AD) cohorts ^53–57^. This is a pattern opposite to that observed acutely after psilocybin in our study sample, of healthy, young participants. Prior work has also highlighted shared miRNA, inflammatory, and neurotrophic alterations across MDD and AD ^58^.

Importantly, immuno-metabolic factors have been implicated in biologically defined subtypes of MDD ^42^, and elevated inflammatory signalling has been reported across psychiatric disorders targeted by psilocybin trials, including depression and anxiety related conditions ^17,18,51^. The IMD concept gives this observation additional clinical context by proposing that systemic inflammation and metabolic disruption define a biologically meaningful subgroup of depression associated with poorer response to standard antidepressant treatment and higher cardiometabolic risk ^42^. Immune activation can also adversely affect synaptic plasticity and BDNF-related mechanisms ^15,41^. Psychedelics or psychedelic-inspired compounds have therefore been proposed as candidates for modulating convergent plasticity and immune pathways in psychiatry ^59–62^. Psilocybin has also been proposed as a potential therapy for AD ^59,62^. This translational interest is supported, although weakly, by a recent case report described transient multidomain functional improvement over subsequent days and weeks in one patient with advanced AD following high-dose psilocybin mushroom administration, but this uncontrolled observation cannot establish disease reversal or disease-modifying efficacy ^63^. Further research in warranted, as psychedelics, or psychedelic-inspired compounds, may hold promise across disorders characterised by altered neuroplasticity and inflammation, spanning psychiatric and neurodegenerative conditions ^15,60,61^.

The present study possesses several strengths. Namely, it is based on a randomized, placebo-controlled, double-blind design with standardized dosing and longitudinal sampling at baseline, +360 minutes post psilocybin dose and +7 days post dose. Small RNAs were profiled using unbiased NGS with rigorous quality control recommendations of the R-ODAF pipeline, including exclusion of haemolytic and low-depth samples and multivariate outlier removal, followed by conservative multi-step statistics which employed elastic net feature selection, covariate adjusted DESeq2 and LMMs, including FDR control. Pathway interpretation relied on experimentally validated miRNA-target interactions, restriction to brain-expressed genes, and cross validation between Reactome and GO to reduce database-driven biases.

However, important limitations constrain interpretation. First, the elastic net analysis was used as an exploratory feature-reduction step rather than within an independent discovery and validation framework. Because DESeq2 and LMM testing were subsequently performed in the same dataset, the resulting statistical evidence is conditional on the preceding feature-selection step. Let-7g-5p and miR-150-5p should therefore be regarded as preliminary candidate signals rather than definitive discoveries controlled for false discovery across the complete analytical pipeline, and independent validation is required. The study cohort consisted of healthy young adults, and it remains to be demonstrated that our findings translate to patient populations or older individuals with inflammation or neurodegeneration. The cohort was also not selected or stratified by depressive symptoms, inflammatory status, metabolic profile or symptoms relevant to AD pathology, limiting inference to IMD or neurodegenerative disease. Second, analyses were performed in blood plasma and infer pathway involvement from circulating miRNA changes rather than from central tissue or cell type specific readouts. We did not experimentally validate the regulatory effects of the identified miRNAs on their candidate target genes, measure miRNA expression in central nervous system tissue, or directly assess synaptic plasticity. Third, ORA and EXOmotif profiling are associative and cannot distinguish primary drug effects from downstream adaptation, selective extracellular vesicle packaging or interindividual differences in baseline biology. Fourth, only two DEmiRs met our predefined significance thresholds, which reduces the risk of false positives but also limits sensitivity to more subtle or inter-individual effects. Fifth, the restricted sampling window of baseline, +360 minutes, and +7 days does not capture the full temporal dynamics of small RNA responses to psilocybin, including earlier peak effects or intermediate changes between the acute and subacute phases. Finally, the primary analyses focused on within-arm longitudinal changes rather than a formal treatment-by-time interaction. The findings should therefore be interpreted as circulating miRNA changes temporally associated with psilocybin exposure, rather than as definitive evidence of psilocybin-specific miRNA regulation. Any extrapolation to AD should also remain cautious, since current human evidence in advanced dementia is limited to anecdotal or early-stage clinical observations, and the recent advanced-AD case report differs substantially from the present study in age, clinical status, dosing form, dose intensity and study design.

In summary, a single low-to-moderate oral dose of psilocybin was associated with a reversible shift in two circulating miRNAs at 360 minutes post dose, which resolved by day 7. Pathway level analysis linked let-7g-5p and miR-150-5p to signalling modules involving neuroplasticity related cascades, cytokine and growth factor pathways, and broad transcriptional control, with sequence features compatible with previous findings regarding extracellular export. The enrichment of cytokine and growth factor pathways further places these findings in the context of emerging immuno-metabolic and mechanism-driven models of depression, although the present healthy-volunteer design does not allow for direct conclusions about clinical subtypes such as IMD. These findings position circulating miRNAs as a feasible and minimally invasive biomarker class for probing acute molecular responses to psilocybin in humans. Future studies should test whether similar miRNA signatures are observed in patient populations, whether they track clinical response and cognitive or affective outcomes, how they relate to extracellular vesicle subtypes, and whether they generalize across different doses, regimens, and psychedelic compounds. Priority should be given to studies integrating miRNA profiling within EV-fractions, and accompanied by inflammatory proteins, metabolic phenotyping, and longitudinal clinical outcomes, particularly in patient populations where immune or metabolic biology may modify treatment response.

## DATA AVAILABILITY

All small RNA-sequencing datasets generated in this study are available in NCBI GEO under accession GSE319559, including raw FASTQ files, processed miRNA count tables, and sample metadata.

## SUPPLEMENTARY MATERIAL

### Let-7g-5p: Reactome first sub-pathways

**Supplementary Table 1:** let-7g-5p First Sub Pathway in order of prevalence - Count & Median FDR, and Percentage based on total across all sub-pathways.

| First sub pathway | Count | Median_FDR | Percentage |
| --- | --- | --- | --- |
| signal transduction | 18 | 0.000302 | 18.18 |
| Generic Transcription Pathway | 9 | 4.44E-06 | 9.09 |
| Insulin Receptor | 6 | 0.000144 | 6.06 |
| MAPK | 6 | 0.000576 | 6.06 |
| Cytokine signalling | 4 | 5.94E-07 | 4.04 |
| Nuclear Receptors | 4 | 0.00078 | 4.04 |
| WNT | 4 | 9.70E-05 | 4.04 |
| Cellular Senescence | 3 | 7.60E-09 | 3.03 |
| Chemical stress | 3 | 0.000864 | 3.03 |
| TGFB | 3 | 2.01E-06 | 3.03 |
| Apoptosis | 2 | 1.70E-05 | 2.02 |
| Melanocyte development | 2 | 0.000462 | 2.02 |
| Non-integrin membrane-ECM interactions | 2 | 0.001243 | 2.02 |
| Post-translational protein modification | 2 | 0.013972 | 2.02 |
| Second messengers | 2 | 0.0017 | 2.02 |
| TP53 Cell death genes | 2 | 0.001161 | 2.02 |
| Cell surface interactions at the vascular wall | 1 | 0.000139 | 1.01 |
| Cellular responses to stimuli | 1 | 1.02E-06 | 1.01 |
| Cellular responses to stress | 1 | 1.59E-06 | 1.01 |
| Death Receptor Signaling | 1 | 0.008526 | 1.01 |
| Degradation of the extracellular matrix | 1 | 0.007237 | 1.01 |
| Developmental Biology | 1 | 0.000393 | 1.01 |
| Disease | 1 | 1.51E-05 | 1.01 |
| ERBB2 | 1 | 0.001623 | 1.01 |
| Extracellular matrix organization | 1 | 0.008526 | 1.01 |
| Gene expression (Transcription) | 1 | 8.46E-09 | 1.01 |
| Hemostasis | 1 | 0.003499 | 1.01 |
| Immune System | 1 | 0.000715 | 1.01 |
| Innate immune system | 1 | 0.011004 | 1.01 |
| Integrin cell surface interactions | 1 | 0.002325 | 1.01 |
| MET | 1 | 0.000275 | 1.01 |
| Mitotic | 1 | 0.008526 | 1.01 |
| PDGF | 1 | 0.00171 | 1.01 |
| Platelet activation, signaling and aggregation | 1 | 0.023102 | 1.01 |
| Programmed Cell Death | 1 | 7.89E-05 | 1.01 |
| Regulation by RUNX4 | 1 | 2.01E-06 | 1.01 |
| Regulation by TP53 | 1 | 0.000381 | 1.01 |
| SCF-KIT | 1 | 0.000849 | 1.01 |
| Signal Transduction | 1 | 1.02E-06 | 1.01 |
| Signaling by NOTCH | 1 | 0.004076 | 1.01 |
| Signaling by Receptor Tyrosine Kinases | 1 | 0.000299 | 1.01 |
| Signaling by VEGF | 1 | 0.006196 | 1.01 |
| VEGF | 1 | 0.004923 | 1.01 |

**Supplementary Table 2:**
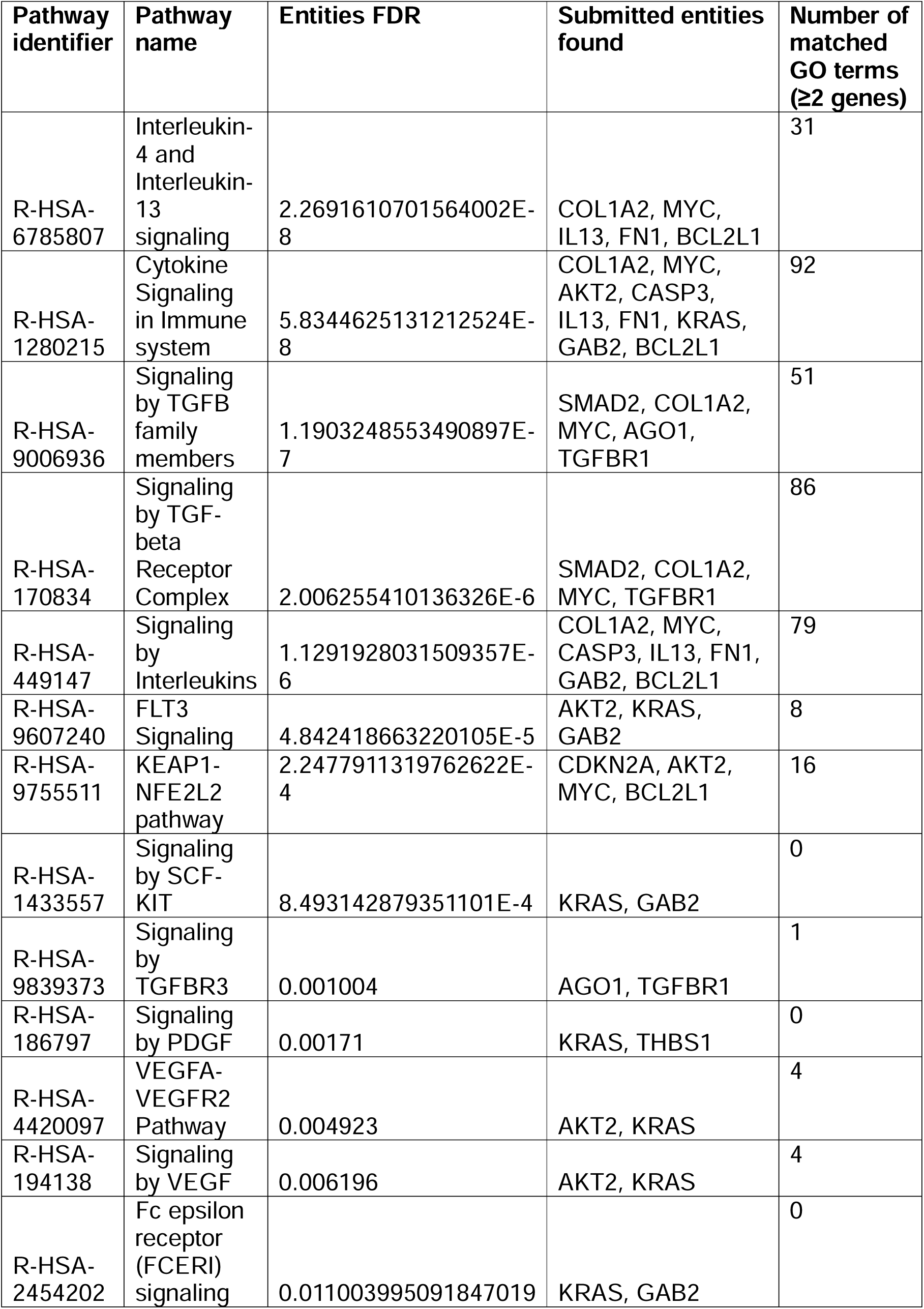
Let-7g-5p: Reactome Cytokine and growth factor signalling pathways.

**Supplementary Table 3:**
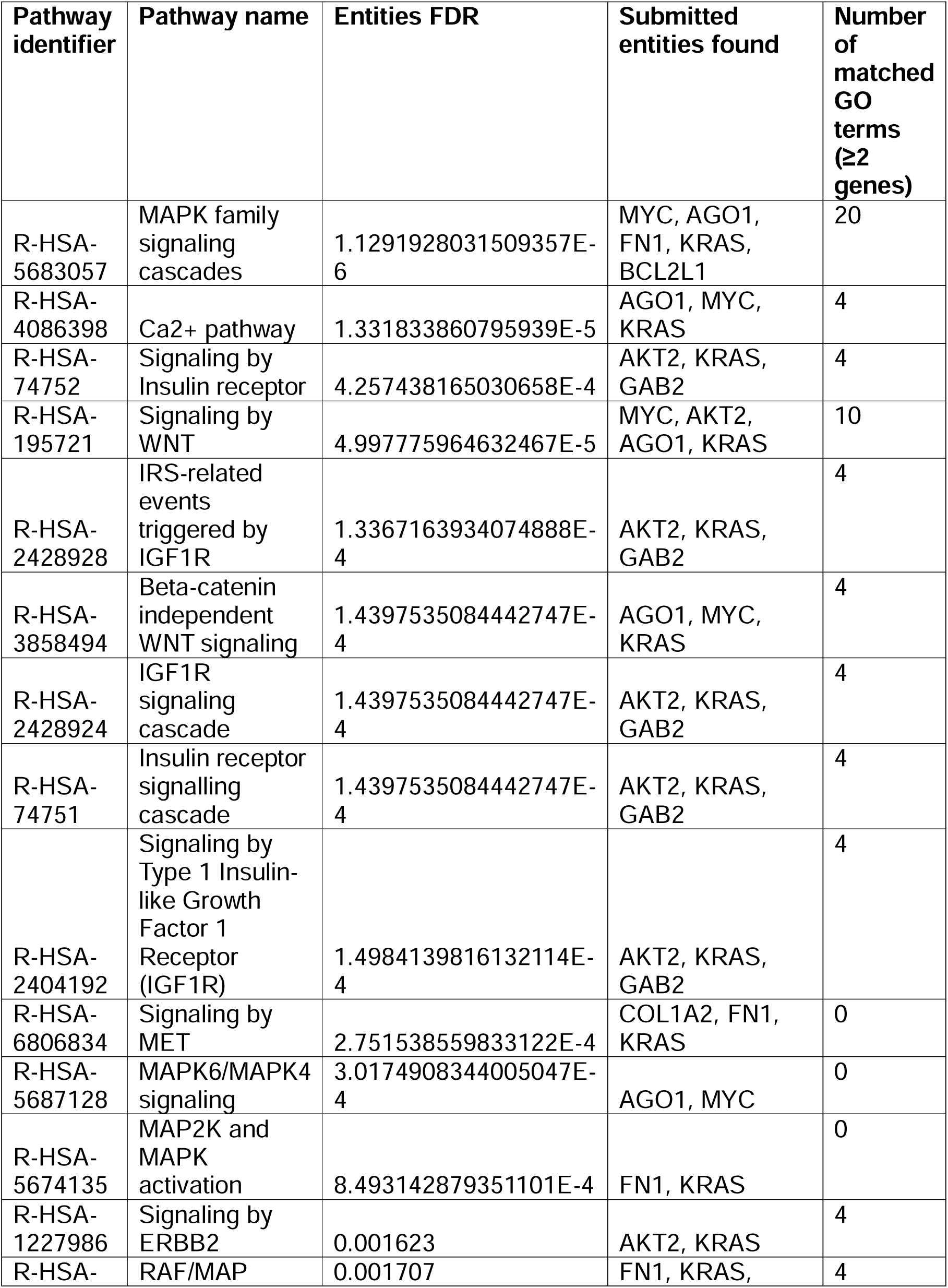
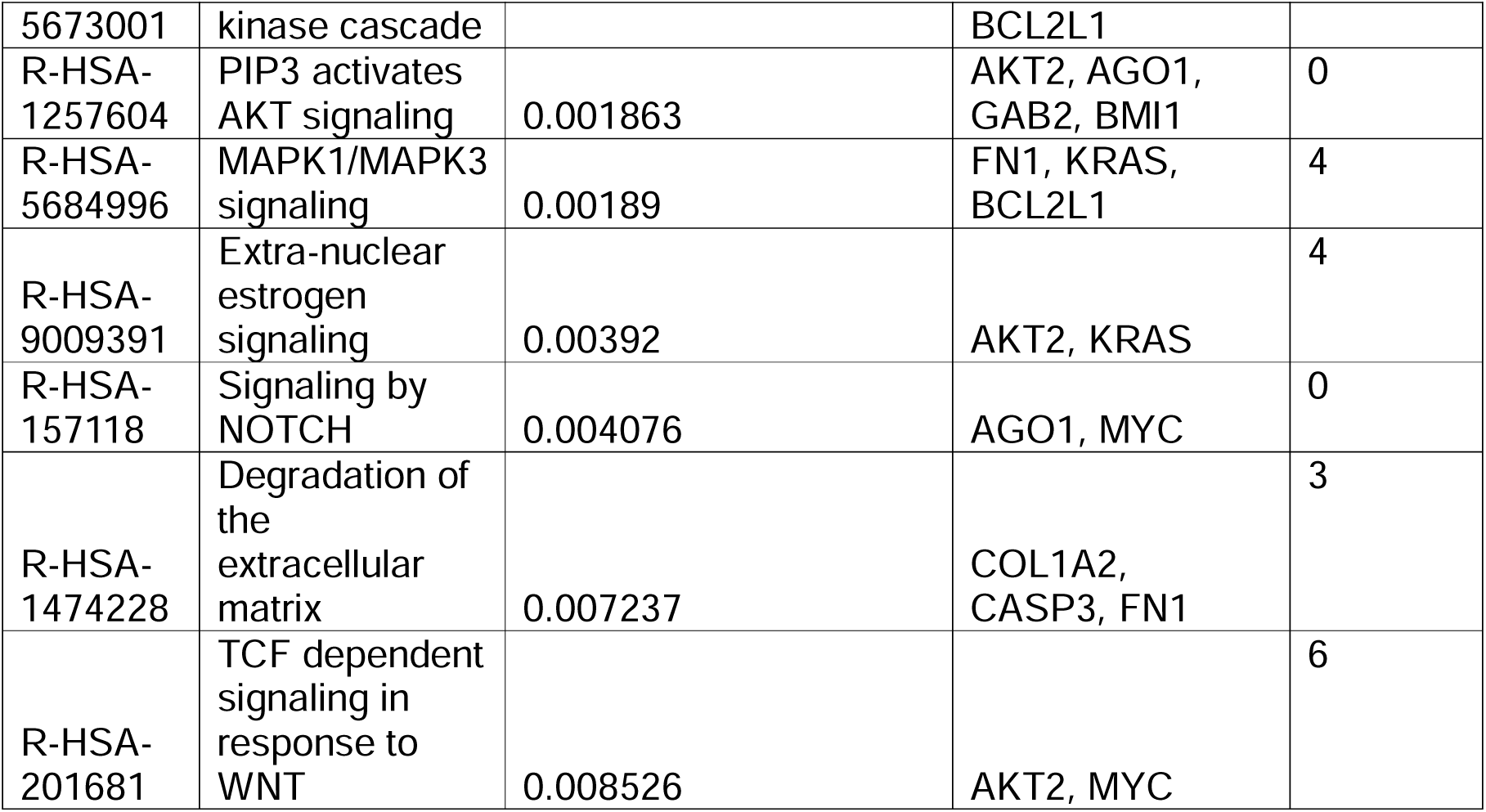
Let-7g-5p: Reactome signal transduction networks.

**Supplementary Table 4:**
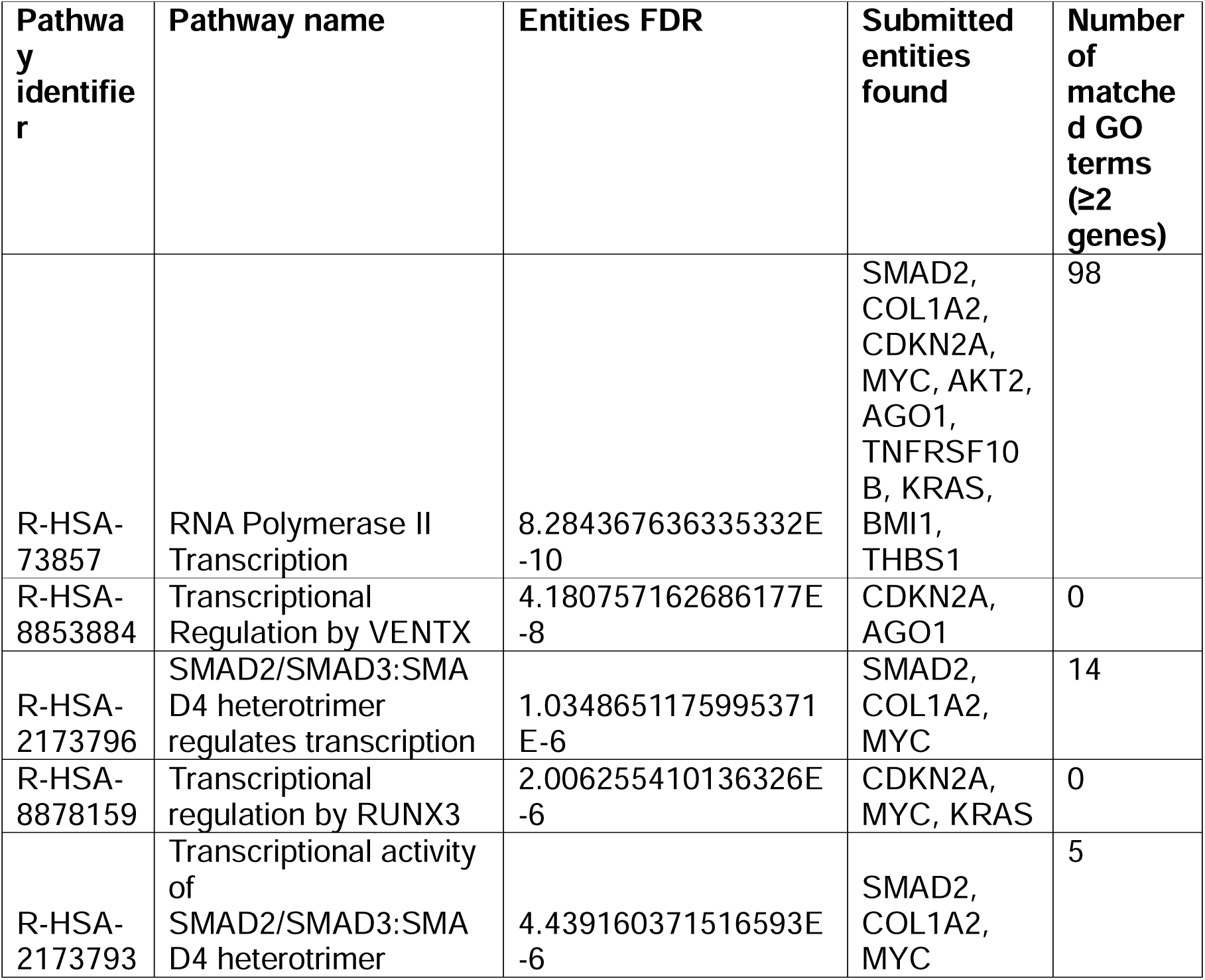
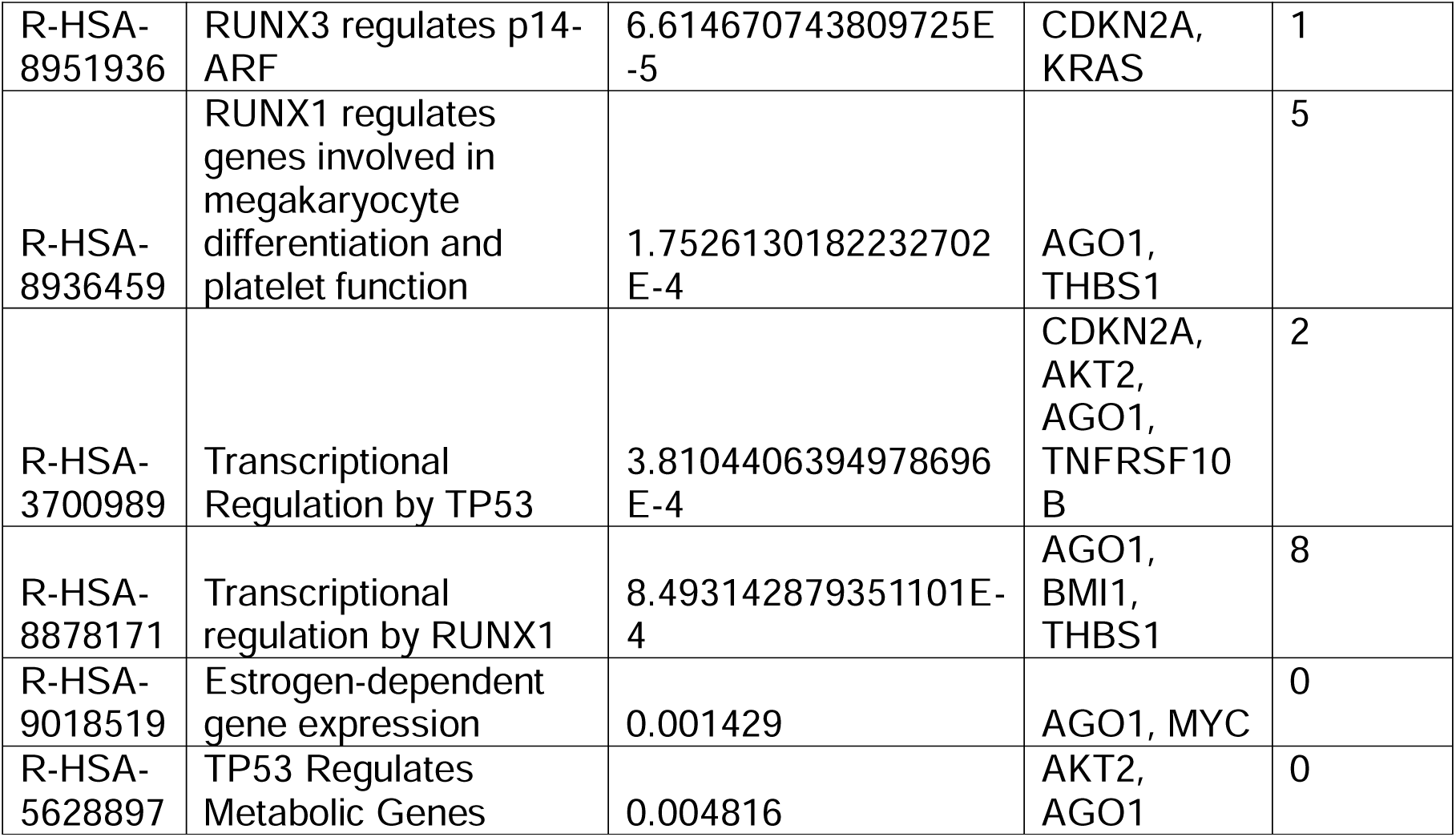
Let-7g-5p: Reactome Gene expression influencing transcription pathways.

### miR-150-5p: Reactome first sub-pathways

**Supplementary Table 5:** miR-150-5p First Sub Pathway in order of prevalence - Count & Median FDR, and Percentage based on total across all sub-pathways.

| First.sub.pathway | Count | Median_FDR | Percentage |
| --- | --- | --- | --- |
| Cytokine signalling | 9 | 0.00263 | 16.07 |
| Generic Transcription Pathway | 4 | 0.001737 | 7.14 |
| NOTCH | 4 | 0.002681 | 7.14 |
| NTRKs | 4 | 0.0083 | 7.14 |
| signal transduction | 4 | 0.002542 | 7.14 |
| Nuclear Receptors | 3 | 0.015662 | 5.36 |
| TGFB | 3 | 0.000946 | 5.36 |
| Hypoxia | 2 | 0.008607 | 3.57 |
| Post-translational protein modification | 2 | 0.02076 | 3.57 |
| Cellular responses to stimuli | 1 | 0.03517 | 1.79 |
| Developmental Biology | 1 | 0.005532 | 1.79 |
| Disease | 1 | 0.0219 | 1.79 |
| GPCR | 1 | 0.011441 | 1.79 |
| Gastrulation | 1 | 0.018288 | 1.79 |
| Gene expression (Transcription) | 1 | 0.002937 | 1.79 |
| Immune System | 1 | 0.000148 | 1.79 |
| Infectious disease | 1 | 0.03517 | 1.79 |
| Maternal to zygotic transition (MZT) | 1 | 0.039604 | 1.79 |
| Mechanical Stimuli | 1 | 0.03517 | 1.79 |
| Melanocyte development | 1 | 0.030384 | 1.79 |
| Nervous system development | 1 | 0.043015 | 1.79 |
| PDGF | 1 | 0.015774 | 1.79 |
| Regulation by AP-2 | 1 | 0.008777 | 1.79 |
| SCF-KIT | 1 | 5.24E-05 | 1.79 |
| Signal Transduction | 1 | 5.75E-06 | 1.79 |
| Signaling by NOTCH | 1 | 8.86E-05 | 1.79 |
| Signaling by Receptor Tyrosine Kinases | 1 | 4.43E-06 | 1.79 |
| Signaling by VEGF | 1 | 0.042003 | 1.79 |
| TP53 Cell death genes | 1 | 0.01969 | 1.79 |
| Transcriptional regulation of granulopoiesis | 1 | 0.015774 | 1.79 |

**Supplementary Table 6:**
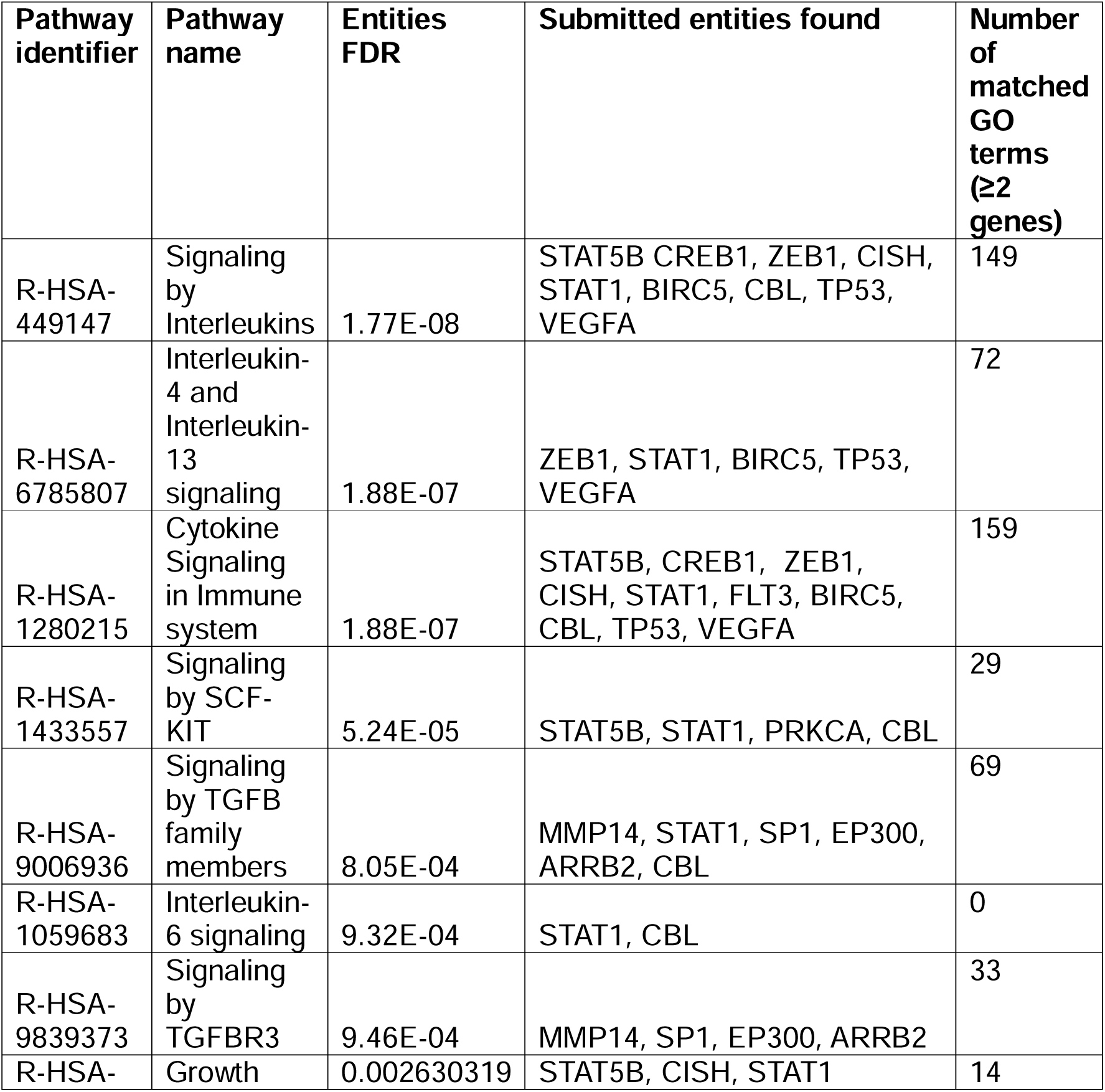
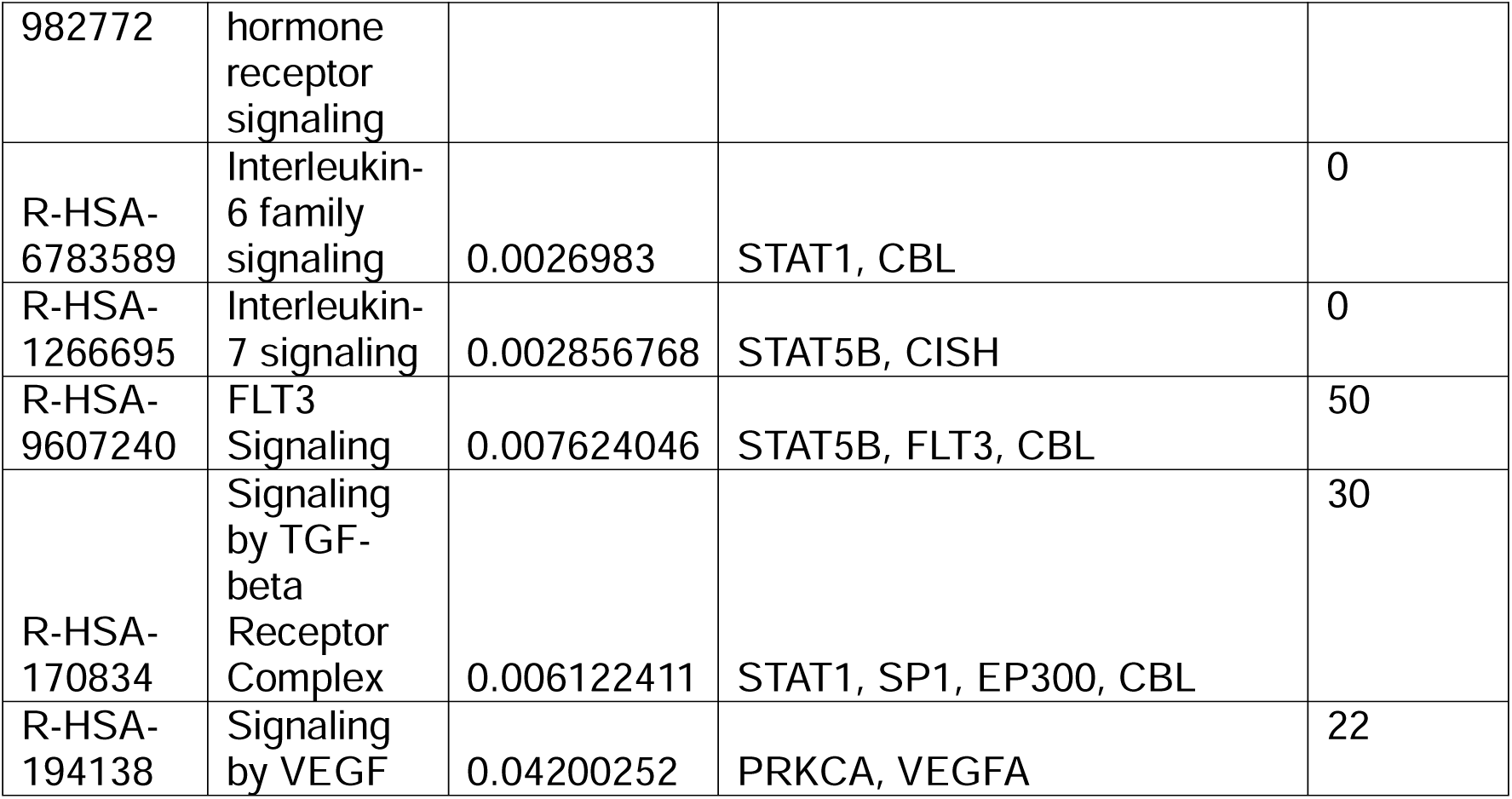
miR-150-5p: Reactome Cytokine and growth factor signalling pathways.

**Supplementary Table 7:**
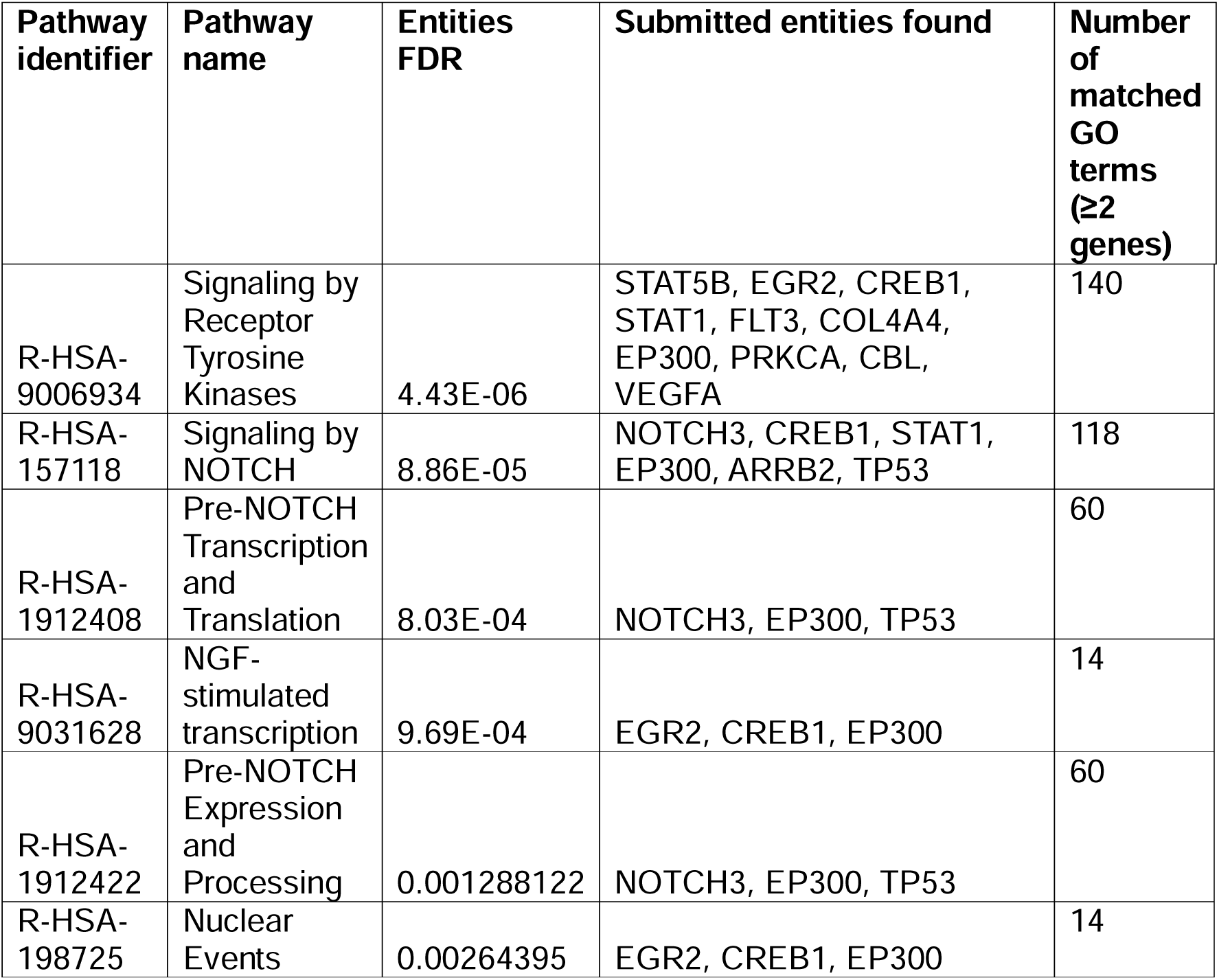
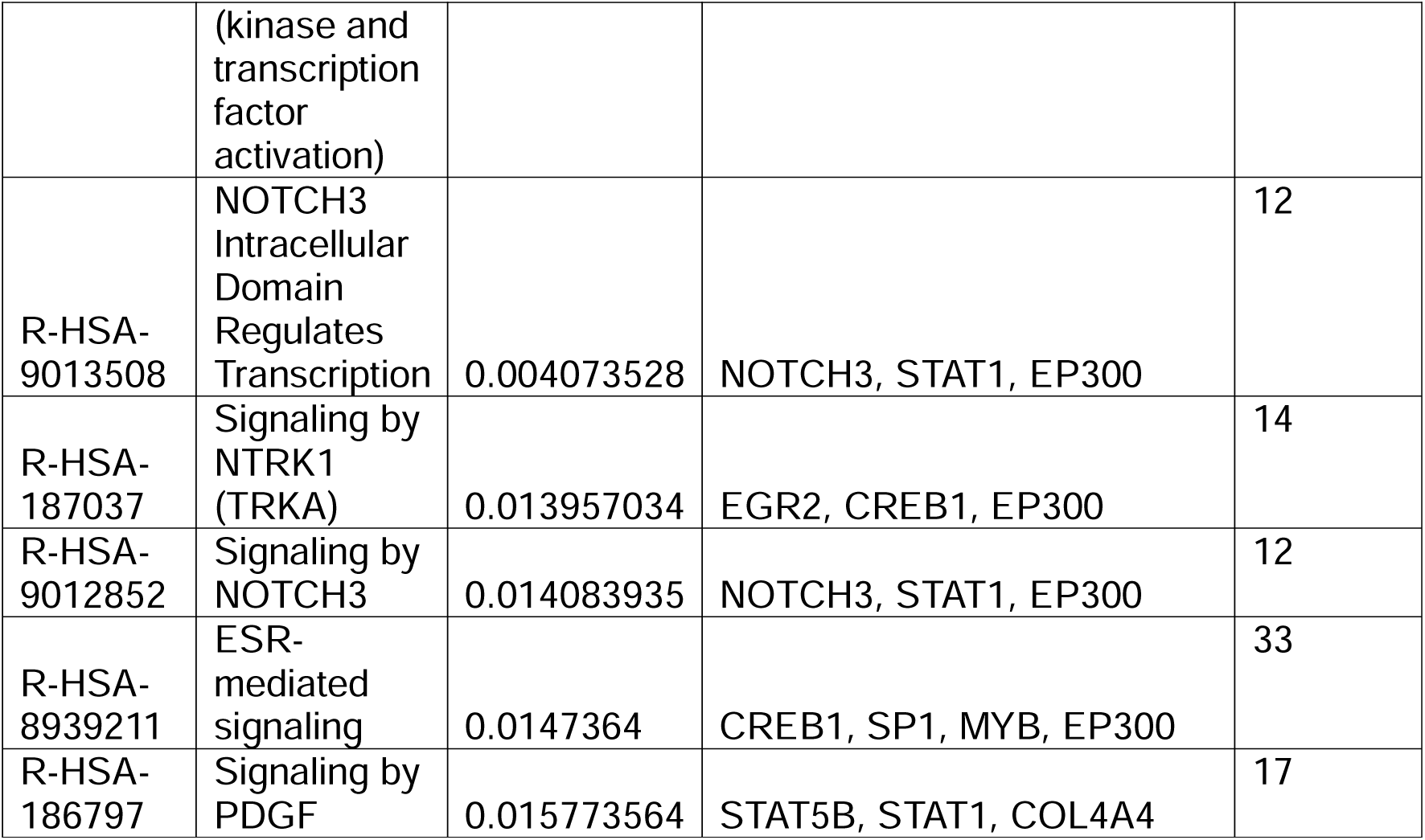
miR-150-5p: Reactome Neurotrophin Signalling and Synaptic Remodelling.

| **Pathway identifier** | **Pathway name** | **Entities FDR** | **Submitted entities found** | **Number of matched GO terms (≥2 genes)** |
| --- | --- | --- | --- | --- |
| R-HSA-9006934 | Signaling by Receptor Tyrosine Kinases | 4.43E-06 | STAT5B, EGR2, CREB1, STAT1, FLT3, COL4A4, EP300, PRKCA, CBL, VEGFA | 140 |
| R-HSA-157118 | Signaling by NOTCH | 8.86E-05 | NOTCH3, CREB1, STAT1, EP300, ARRB2, TP53 | 118 |
| R-HSA-1912408 | Pre-NOTCH Transcription and Translation | 8.03E-04 | NOTCH3, EP300, TP53 | 60 |
| R-HSA-9031628 | NGF-stimulated transcription | 9.69E-04 | EGR2, CREB1, EP300 | 14 |
| R-HSA-1912422 | Pre-NOTCH Expression and Processing | 0.001288122 | NOTCH3, EP300, TP53 | 60 |
| R-HSA-198725 | Nuclear Events (kinase and transcription factor activation) | 0.00264395 | EGR2, CREB1, EP300 | 14 |
| R-HSA-9013508 | NOTCH3 Intracellular Domain Regulates Transcription | 0.004073528 | NOTCH3, STAT1, EP300 | 12 |
| R-HSA-187037 | Signaling by NTRK1 (TRKA) | 0.013957034 | EGR2, CREB1, EP300 | 14 |
| R-HSA-9012852 | Signaling by NOTCH3 | 0.014083935 | NOTCH3, STAT1, EP300 | 12 |
| R-HSA-8939211 | ESR-mediated signaling | 0.0147364 | CREB1, SP1, MYB, EP300 | 33 |
| R-HSA-186797 | Signaling by PDGF | 0.015773564 | STAT5B, STAT1, COL4A4 | 17 |

**Supplementary Table 8:**
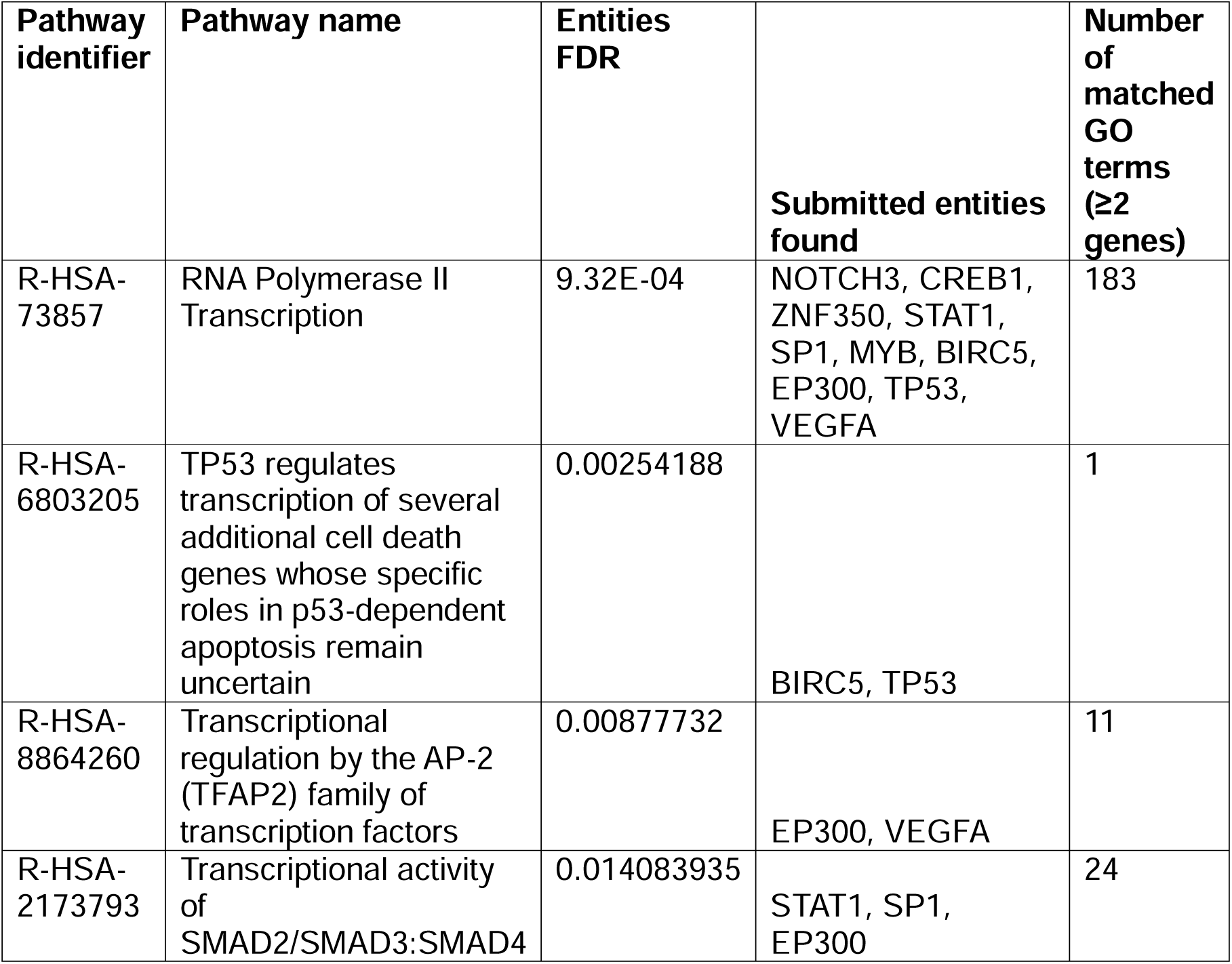
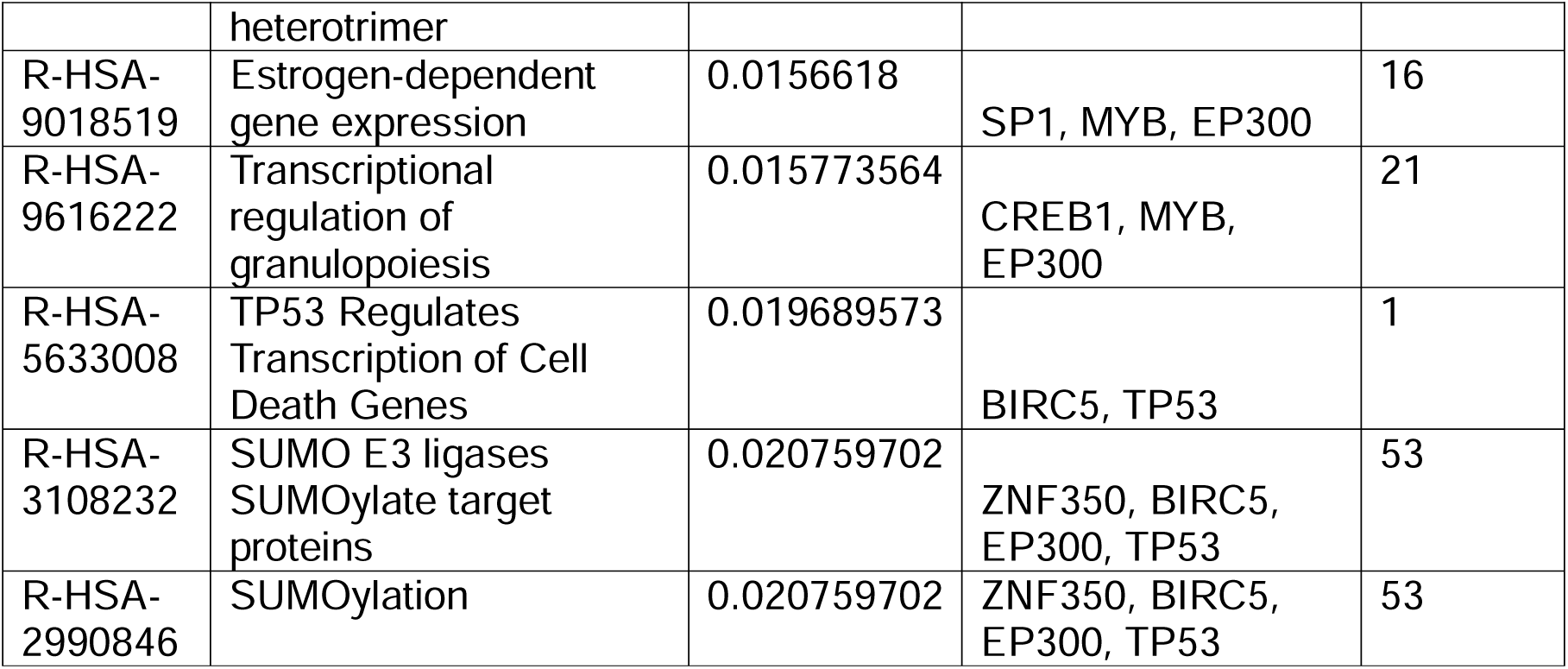
miR-150-5p: Reactome Gene expression influencing transcription pathways.

### Supplementary 8 Unique Reactome pathways with 2 or more genes overlapping with GO terms.

Unique to hsa-let-7g-5p

- MAPK family signaling cascades
- Transcriptional regulation by RUNX3
- RUNX3 regulates p14-ARF
- IGF1R signaling cascade
- RUNX1 regulates genes involved in megakaryocyte differentiation and platelet function
- Signaling by TGF-beta Receptor Complex in Cancer
- Transcriptional regulation by RUNX1

Unique to hsa-mir-150-5p

- Signaling by NOTCH
- NGF-stimulated transcription
- Signaling by NTRK1 (TRKA)
- Transcriptional regulation by the AP-2 (TFAP2) family of transcription factors
- Signaling by NOTCH3
- Signaling by PDGF
- SUMOylation
- Signaling by NOTCH2

Common to both

- Cytokine Signaling in Immune system
- RNA Polymerase II Transcription
- Signaling by Interleukins
- Interleukin-4 and Interleukin-13 signaling
- Signaling by TGF-beta Receptor Complex
- Signaling by Receptor Tyrosine Kinases
- SMAD2/SMAD3:SMAD4 heterotrimer regulates transcription
- Transcriptional activity of SMAD2/SMAD3:SMAD4 heterotrimer
- Signaling by VEGF

**Supplementary 9.** Age/Sex/BMI significance.

| Parameter | miRNA | baseMean | log2FoldChange | lfcSE | stat | pvalue | padj |
| --- | --- | --- | --- | --- | --- | --- | --- |
| Sex | hsa-miR-106b-3p | 1049.387 | 0.231208 | 0.049697 | 4.652402 | 3.28E-06 | 0.000274 |
| Sex | hsa-miR-132-3p | 123.6571 | 0.192325 | 0.067694 | 2.841085 | 0.004496 | 0.080833 |
| Sex | hsa-miR-154-3p | 43.51604 | -0.5562 | 0.168379 | -3.30324 | 0.000956 | 0.026601 |
| Sex | hsa-miR-190a-5p | 25.0322 | -0.43741 | 0.132073 | -3.31188 | 0.000927 | NA |
| Sex | hsa-miR-4286 | 801.9613 | 0.362127 | 0.076519 | 4.732535 | 2.22E-06 | 0.000247 |
| Sex | hsa-miR-454-5p | 70.26866 | 0.264402 | 0.075051 | 3.522989 | 0.000427 | 0.014252 |
| Sex | hsa-miR-4742-3p | 16.24272 | 0.299109 | 0.131503 | 2.274549 | 0.022933 | NA |
| Sex | hsa-miR-493-5p | 1708.576 | -0.31854 | 0.130067 | -2.44903 | 0.014324 | 0.144978 |
| Age | hsa-miR-5187-5p | 35.38088 | 0.66242 | 0.204798 | 3.234505 | 0.001219 | 0.149351 |
| BMI | hsa-miR-1-3p | 1423.112 | -0.77916 | 0.17404 | -4.47692 | 7.57E-06 | 0.003082 |
| BMI | hsa-miR-151a-5p/151b | 20029.38 | -0.1615 | 0.055269 | -2.92216 | 0.003476 | 0.09985 |
| BMI | hsa-miR-26b-5p | 24201.15 | -0.27071 | 0.064117 | -4.22216 | 2.42E-05 | 0.004924 |
| BMI | hsa-miR-6859-5p | 20.94783 | 0.297681 | 0.140082 | 2.125057 | 0.033582 | NA |

### Supplementary 10

**Gene Targets of let-7g-5p:**

AGO1, AKT2, BCL2L1, BMI1, CASP3, CDKN2A, COL1A2, FN1, GAB2, HMGA2, IL13, KRAS, MYC, SMAD2, TGFBR1, THBS1, TNFRSF10B.

**Gene Targets of miR-150-5p:**

ARRB2, BIRC5, CBL, CCR6, CISH, COL4A4, CREB1, CXCR4, EP300, EGR2, FLT3, MMP14, MYB, NOTCH3, P2RX7, POLD3, PRKCA, SLC2A1, SP1, SRCIN1, STAT1, STAT5B, TP53, VEGFA, ZEB1, ZNF350, ZNRD2

